# Unbiased Population-Based Statistics to Obtain Pathologic Burden of Injury after Experimental TBI

**DOI:** 10.1101/2025.04.03.647083

**Authors:** G. Smith, C. Santana-Gomez, R.J. Staba, N.G. Harris

**Affiliations:** UCLA Brain Injury Research Center, Department of Neurosurgery, Geffen Medical School; Intellectual Development and Disabilities Research Center; Department of Neurology, University of California at Los Angeles, Los Angeles, CA, 90095, USA

**Keywords:** Traumatic Brain Injury, rigor, DWI, diffusion imaging

## Abstract

Reproducibility of scientific data is a current concern throughout the neuroscience field. There are multiple on-going efforts to help resolve this problem. Within the preclinical neuroimaging field, the continued use of a region-of interest (ROI) type approaches combined with the well-known spatial heterogeneity of traumatic brain injury pathology is a barrier to the replicability and repeatability of data. Here we propose the conjoint use of an unbiased analysis of the whole brain after injury together with a population-based statistical analysis of sham-control brains as one approach that has been used in clinical research to help resolve this issue. The approach produces two volumes of pathology that are outside the normal range of sham brains, and can be interpreted as whole brain burden of injury. Using diffusion weighted imaging derived scalars from a tensor analysis of data acquired from adult, male rats at 2, 9 days, 1 and 5 months after lateral fluid percussion injury (LFPI) and in shams (n=73 and 12, respectively), we compared a data-driven, z-score mapping method to a whole brain and white matter-specific analysis, as well as an ROI-based analysis with brain regions preselected by virtue of their large group effect sizes. We show that the data-driven approach is statistically robust, providing the advantage of a large group effect size typical of a ROI analysis of mean scalar values derived from the tensor in regions of gross injury, but without the large multi-region statistical correction required for interrogating multiple brain areas, and without the potential bias inherent with using preselected ROIs. We show that the technique correctly captures the expected longitudinal time-course of the diffusion scalar volumes based on the spatial extent of the pathology and the known temporal changes in scalar values in the LFPI model.

## Introduction

Reproducibility of data is a current, major problem in preclinical scientific research(Begley and Ioannidis, 2015; Prinz et al., 2011; “Trouble at the lab,” 2013) that is presumably associated with poor clinical translation in the field of traumatic brain injury(Maas et al., 2017; Stein, 2015). There are plans to address this(Collins and Tabak, 2014) as led by the Findability, Accessibility, Interoperability and Reuse (FAIR) guiding principles(Amadi et al., 2024) developed to help solve problems of reproducibility in research, along with the Animal Research: Reporting of In Vivo Experiments (ARRIVE) guidelines(Sert et al., 2020), and the increasingly used common data elements for data reporting and sharing in the traumatic brain injury (TBI) field (LaPlaca et al., 2021; Smith et al., 2015).

Within neuroimaging research, the classical region-of-interest (ROI)-based analysis approach has often been used in the TBI research field because it provides an effective way to clearly delineate group differences. However, one issue with this approach is that regions are not always chosen or located in a standardized way relative to a brain atlas or to landmark features. Even given the implementation of co-registration methods to enable the use of systematic, parcellated atlases, ROI-derived data may not be sensitive enough to produce significant group effect sizes given the well-known spatial heterogeneous effects of TBI. One method often used for ROI-based analysis of the injured brain includes a-priori justification for use of specific ROIs. This is typically implemented using hypothesis-driven information available on specific brain regions targeted for analysis, or other methods of pre-selecting ROIs, such as using peri-contused regions in those cases where a primary injured site or focal injury is present. However, this approach is not useful when modelling the majority of brain injuries which are milder, and present with little to no obvious gross pathology, and this precludes the preselection of ROI targets of interest. Furthermore, this predetermined selection of targets may also be biased against unexpected pathology that may be missed by focusing on brain regions typically analyzed within the field.

As an alternative, a tract-based statistical analysis is a popular approach deployed within the field that is unbiased to white matter region( Harris et al., 2016; Wright et al., 2017). However, it requires good overlap of pathology between subjects to arrive at a significant effect size. Whole brain, voxel-wise group comparisons can be a useful tool for grading the pathologic severity of a TBI in a more reproducibility manner and with a focus on the unbiased nature of the method. The advantage of this approach is that not only is it regionally unbiased, but that it retains the spatial complexity that a multi-level, ROI interrogation would offer through use of voxel-level information. However, the statistical correction required for the multivoxel comparison reduces the power to detect a group difference in much the same way that a parcellated atlas, ROI-based approach does, potentially missing data of interest for differentiating a group effect. These technical, image processing steps that prevent the distillation of complex pathology to a single, simplified value also slow the adoption of imaging biomarkers into the creation of statistical models to permit inferences on injury severity or effect of intervention.

Here, we investigate a method to address the inherent problems of these two orthogonal approaches through the conjoint use of an unbiased analysis of injured brain together with a population-based statistical analysis of sham-control brains. Variations of this approach have been applied to clinical data analysis over the last decade or so(Davenport et al., 2012; Jorge et al., 2012; Kim et al., 2013; Ling et al., 2012; Lipton et al., 2012; Mayer et al., 2014, 2012; Watts et al., 2014), yet have not been applied to preclinical work. This entirely data-driven approach is conducted in a common 3-dimensional space so that the spatial information encoded within each brain is ultimately used to create z-score maps from which to statistically delineate boundaries of pathology within each injured brain. The result is the delineation of two volumes of tissue per brain that lie above or below the normal range of sham control brain values. This burden of pathology, or volumes of interest that is described by these single values can then be used with other parameters for further statistical analysis of prediction of outcome or intervention. Recently the critical importance of the statistical correction of the resultant data has reemerged(Gimbel et al., 2024a; Mayer et al., 2024), emphasizing the necessity of correcting for potential bias. Here we employ statistical theory-derived distribution-corrected z-score (DisCo-Z) adjustments to the delineation of pathologic tissue in injured brain compared to a reference sham group, first proposed by Mayer and colleagues for the analysis of clinical TBI data(Mayer et al., 2014).

In the current work we used diffusion weighted imaging derived scalars from a tensor analysis of data acquired at 2d to 5m after lateral fluid percussion injury (LFPI) in adult rats. We compared a data-driven, z-score mapping method to a whole brain and white matter-specific analysis, as well as an ROI-based analysis. We also preselected ROIs based on the location of the primary injury for potential maximum effect size, and used regions immediately adjacent and ventral to the craniectomy and the corresponding contralateral regions. We show that the data-driven approach provides many of the advantages of a typical ROI analysis but without the large statistical correction required for interrogating multiple regions, and without the potential bias inherent in preselected ROIs.

## Methods

### Experimental Paradigm

Diffusion weighted imaging data were acquired from adult, Sprague Dawley rats (318±36.6g body weight at time of injury) at 2d, 9d, 1m and 5m after moderate-severe, LFPI and from sham-craniectomy controls. A total of 85 animals were randomized to sham (n=12) or TBI groups (n=73). Data were acquired in cohorts of 8 rats consisting of 7 injured and 1 sham. From the TBI group, 31 never recovered from apnea at the time of injury, and 5 were excluded due to a dura tear at the time of injury. MRI data were acquired from the resulting sham and TBI rats (n=12 and 37, respectively) at 2d, 9d, 1m, 5m post-injury. There was a further loss of data due to death during MRI scanning (n=2 TBI) or unrelated to scanning (n=1 TBI), and due to scan failure as a result of hardware problems in 9 rats (10 scan failures in both sham and TBI). As a result, temporally complete data sets were acquired at all 4 timepoints from 8 sham and 24 TBI rats, but scans from incomplete datasets were also included, resulting in the final groupXtime size of n=10,10,11,12 for sham and n=36,32,33,33 for TBI rats at 2d, 9d, 1m, 5m post-injury, respectively. These data were collected as part of the EpiBioS4Rx consortium and the metadata from these including signal-to-noise, acquisition parameters and physiology data have been partly published(Immonen et al., 2019a, 2019b). All study protocols used complied with the Public Health Service Policy on Humane Care and Use of Laboratory Animals and were approved by the Chancellor’s Animal Research Committee, University of California, Los Angeles, California.

### Lateral Fluid Percussion Injury

LFP was conducted as before (Fox et al., 2024; Santana-Gomez et al., 2024). Briefly, while under general anesthesia (2% isoflurane vaporized in oxygen flowing at 0.8 l/min; 5% induction for 30.2±5.9 minutes, the skin prepared for aseptic surgery, a midline incision was made, and rats received a 5mm craniectomy using a manual trephine (centered at -4.5mm to bregma and 2.5mm left of the midline). A moderate to severe fluid percussion injury was then delivered using 19-23° angle to generate 2.2±0.2 atmosphere pressure. The duration of post-injury apnea was recorded (24.9±18.2 seconds). For the sham controls only the craniectomy was performed and using the same duration of anesthesia. The skin was closed with sutures and the rat recovered.

### MRI Acquisition

Diffusion weighted imaging (DWI) data were acquired from all rats under 1.5% isoflurane anesthesia vaporized in room air and using a 7T Bruker MRI at 2 and 9 days, and then 1 and 5m post injury. Images were acquired with a 3-dimensional spin echo, single shot, echo planar sequence constrained to a data matrix of 96×72×49 (readout, phase-encode1, phase-encode2) using lateral, dorsal and ventral outer-volume signal suppression saturation slices, resulting in a 250mm^3^ isotropic resolution. Four B0s and 42 gradient vectors at a b-value of 2800 s/mm^2^ were acquired with a TR/TE of 1000/26ms, and a diffusion time of 12ms resulting in a total acquisition time of 37 mins. These data were recorded following a number of other sequences run as part of a larger study, so that the total time in the magnet was 110 mins/session.

### Image Preprocessing

Preprocessing for MRI began with converting the Bruker data to NifTi using Brkraw (Lee et al., 2020) followed by a pipeline implemented in MRTRIX(Tournier et al., 2019) with FSL tools(Jenkinson et al., 2012; Smith et al., 2004; Woolrich et al., 2009). Data were brain extracted(Smith, 2002), denoised(Cordero-Grande et al., 2019; Veraart et al., 2016), corrected for Gibbs ringing(Kellner et al., 2016), and then corrected for geometric distortions due to gradient-induced eddy current fields and susceptibility-induced off-resonance field using TOPUP(Andersson et al., 2003) and Eddy(Andersson et al., 2016; Andersson and Sotiropoulos, 2016). After fitting the data to the tensor, preliminary fractional anisotropy (FA) maps were created for each animal in subject space, and an unbiased mean deformation template (MDT) was built using rigid, affine, and non-linear alignment registrations(Avants et al., 2008). The quality of the image co-registrations were manually checked for each data set. In cases of severe brain injury with large cavity formations and thinned and/or displaced corpus callosum, attempts were made to use lesion/cavity masking to weight the registration toward intact areas. All scans, regardless of whether they contained small, acquisition-specific artifacts were entered into the analysis. Those scans that contained whole-brain artifacts related to scan problems were noted as scan failures and were removed from the study, as noted in the experimental paradigm section above.

### Z-score Analysis

Subject-specific, tensor-derived scalars (FA, radial, axial and mean diffusivity-RD, AD, MD, respectively) were transformed to common, MDT space using the generated warp fields and affine transformations created from the FA based MDT formation. The voxel-wise mean and standard deviation were calculated for the sham rats for each scalar at each time point, which were then used to compute scalar-specific, voxel-wise z-value maps, for all injured and sham subject data. Data were then thresholded using a distribution-corrected z-score (DisCo-Z)(Mayer et al., 2014) based on a desired z value of |z|>±2.33 (p<0.05, **Fig. 1**). The resulting z value for thresholding the injury groups at 2d, 9d, 1m, and 5m post injury were, 2.96, 2.96, 2.84, and 2.79 respectively. For the sham group the adjusted z values were 2.04, 2.04, 2.07, and 2.09 for their respective timepoint. The brain-wide volumes of tissue surviving these upper and lower thresholds for each tensor scalar, herein designated Scalar_LOW_ and Scalar_HIGH_ volumes, were then quantified for each brain, and for each z threshold above and below the sham mean values. Group-level maps of each Scalar_LOW_ and Scalar_HIGH_ volumes are shown as incidence maps showing the overlap within each group. For the average scalar value analysis of these data, a single mask derived from all animals at each timepoint was used so that the same volume of tissue was compared across subjects. This mask was constructed from brain voxels in which greater than 10% of all injured rats were significantly different from shams.

**Fig. 1.**
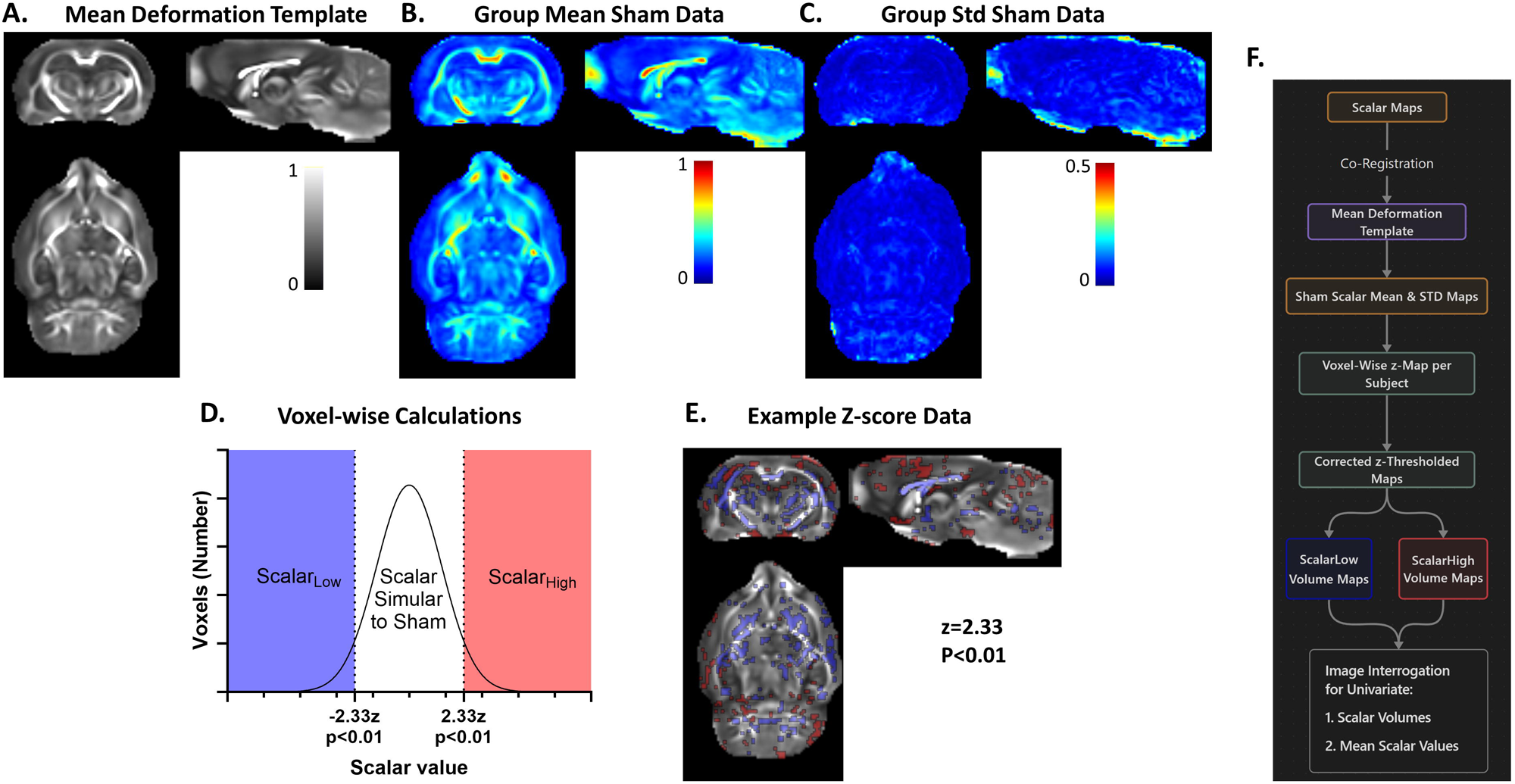
Overview of Z-map creation for FA. **[A]** After creating a mean deformation template (MDT) from all available sham and injury FA data within the sample population, sham group [**B**] mean and [**C**] standard deviation maps are calculated from the MDT co-registered data for each DWI tensor scalar (FA, MD, RD, AD) which are then used to construct [**D**] voxel-wise z-score thresholds at ±2.33z (P<0.05) for each subject to enable the computation of volumes of pathology composed of Scalar values higher or lower than this threshold (Scalar_High_ and Scalar_Low_, respectively) indicated by the red and purple regions respectively. [**E**] Representative injured rat showing the brain voxels that are significantly higher or lower (red and blue, respectively) compared to the sham population. [**F**] Summary overview of the analysis pipeline: from scalar map input to interrogation of the images for scalar volume above or below the corrected z-threshold.

### ROI-based analysis

A parcellated rodent brain atlas (Harris et al., 2016; Valdés-hernández et al., 2011) was co-registered into MDT template space for region-of-interest (ROI)-based analysis. Regions corresponding to the hippocampus and thalamus subdivided into left, right, anterior and posterior regions, as well as the left and right: sensory and motor cortices, striatum and genu of the corpus callosum were used for analysis.

### Statistics

All data were displayed as violin plots showing medians and quartile ranges. Group-level analysis of z-score and ROI-derived volume data were non-normally distributed and so group differences in Scalar_LOW_ and Scalar_HIGH_ volumes and values at 1m post-injury volume were tested for differences using the Kruskal-Wallis test with Dunn’s correction for multiple ROI comparisons. For the group difference in mean FA between injury and sham animals within each ROI, the Mann-Whitney U test was utilized. Group effect size was calculated using Hedges bias-corrected effect size. Tensor scalars from the same brains were treated as independent measures given the preponderance of literature evidence across multiple disease states that they demarcate different pathological processes. In addition, given the spatially discrete distributions of the Scalar_LOW_ and Scalar_HIGH_ sample populations throughout the results that likely indicate dissimilar pathologic processes, we also elected to treat these as independent variables. *<u>Scalar Values</u>:* For the temporal analysis of the mean scalar values, the 3 ROIs with the 3 strongest effect sizes along with the whole brain and whole brain white matter tissue were selected for comparison to the data-driven, tissue analysis. Temporal analysis of mean scalar values within injured rats was conducted using a 1-way, mixed-effects analysis of variance to test for the effect of time after injury using Tukey’s multiple comparison correction. *<u>Scalar Volumes</u>:* For group comparison of Scalar_LOW_ and Scalar_HIGH_ volumes at each post-injury time point, 2-way repeated measures, mixed-effects model ANOVA with Sidak’s test for multiple comparison was applied for each scalar. To determine change in Scalar_LOW_ and Scalar_HIGH_ volumes over time within the injury group, a 1-way, mixed-effects analysis of variance was used together with Sidaks’s test for multiple comparison.

## Results

We first compared the two image interrogation techniques: the data-driven z-score mapping and ROI-base analyses. We limited the analysis to 1mo post-injury data because the diffusivity properties of the injured tissue are more stable at this post-injury time-point than earlier.

### Data-Driven Analysis of Whole Brain Injury Burden: Volumes of Pathology

Application of the data-driven, voxel-based method (z=2.33, P<.01; **Fig. 1**) to whole brain FA images derived from DWI data acquired from a group of sham and LFPI rats 1mo after injury revealed a significant increase in volume of brain pathology after injury. This increase was composed of both higher and lower FA values (FA_High_, FA_Low_ volumes, P<.05, P<.001, respectively) when compared to volumes derived from the sham population (**Fig. 2A**). The spatial distribution of these pathologic volumes of tissue was best exhibited by voxel overlap maps that describe the incidence of voxels in each brain that survived the FA_High_ and FA_Low_ z-score thresholds. This revealed the expected bias towards the primary, left hemispheric injury (**Fig. 2B**). The peak overlap of FA_Low_ volumes among 33 injured rats occurred within the ipsilateral corpus callosum, with further overlap among 50-75% of rats in contralateral corpus callosum, ipsilateral fimbria and internal capsule, as well as ipsilateral cortical grey matter (**Fig. 2B**). The peak overlap of FA_High_ volumes was not as consistent within the LFPI group compared to FA_Low_, with only around 50% overlap between pathologic-identified regions, and that was constrained to the ipsilateral, deep cortical grey matter immediately adjacent to regions of grey matter FA_Low_, and to some bilateral, dorsal hippocampal regions. These FA_High_ and FA_Low_ volume data resulted in an effect size of 0.05 and 1.43, respectively to discriminate between sham and injured groups, and this achieved a statistical power of 0.97 for the FA_Low_ tissue volume. We found similar results for the other tensor scalars: mean, axial and radial diffusivity (**S1-3**), although the effect size was greatest for FA_Low_.

**Fig. 2.**
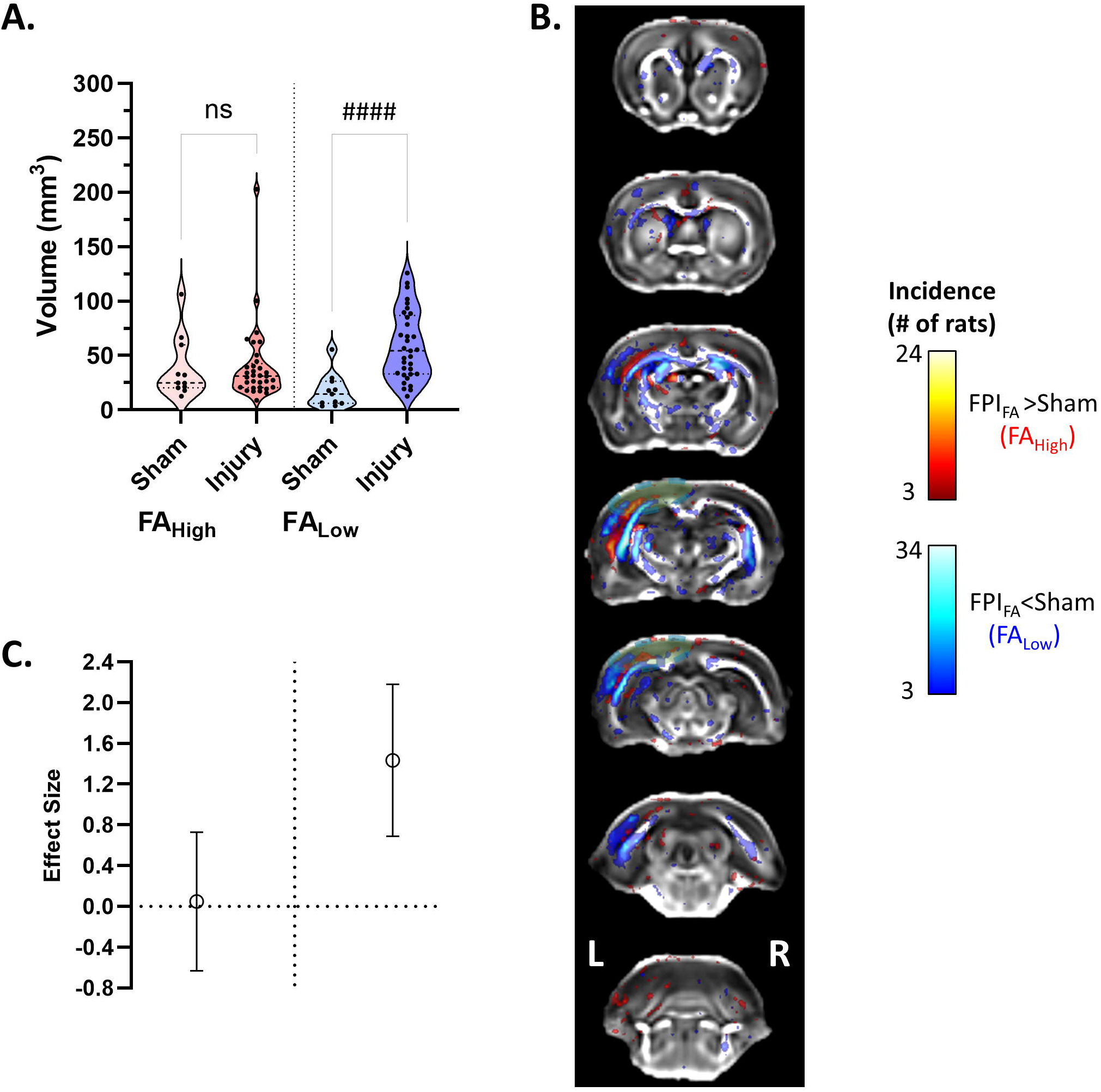
Data-driven Analysis of FA at 1month after injury. [**A]** Volumes of FA_High_ and FA_Low_ in sham and LFPI rats. [**B]** A population incidence map of FA_High_ and FA_Low_ volumes from all injured rats highlighting the location of altered tissue (voxel pseudocolors indicate number of rats). [**C]** Graph of statistical effect size, (Hedges’ g) with confidence intervals for the effect of group for FA_High_ and FA_Low_ volumes. Kruskal-Wallis test #### p<0.0001. Green disk represents the location of craniectomy.

### Comparison of Data-Driven versus ROI Approach to Diffusion Scalar Value Analysis

We compared the data-driven approach for assessing group FA value differences to using large ROIs encompassing whole brain grey and white matter, and separately from white matter at 1mo post-LFPI (**Fig. 3A**). The data-driven FA values were obtained from the mean of the voxel values that composed each FA_High_ and FA_Low_-derived volumes. While the mean FA values in the two large ROIs were significantly lower in injured compared to sham brains (Mann-Whitney U=88, P<.05:U=69, P<0.01 respectively, uncorrected for multiple comparisons due to this head-to-head comparison), the difference was more significant using the data-driven method to identify pathologic tissue (FA_High_ and FA_Low_) using masks constrained to voxels containing >10% overlap of rats in each group (Mann-Whitney U=75, P<0.01: U=11, P<.0001, respectively, uncorrected **Fig. 3A**). This effect was aided by lower within-group variability of data obtained through the data-driven method, compared to the large ROI-based approach. This culminated in a greater effect size between injured and sham groups for the data-driven approach (-2.14, 0.92 for FA_Low_ and FA_High_, respectively) compared to the whole brain white matter and white+grey matter large ROIs (-1.03 and -1.13 respectively, **Fig. 3B**). We found similar results for the other tensor scalars: mean, axial and radial diffusivity (**S4-6**), and once again the effect size was greatest for FA_Low_.

**Fig. 3.**
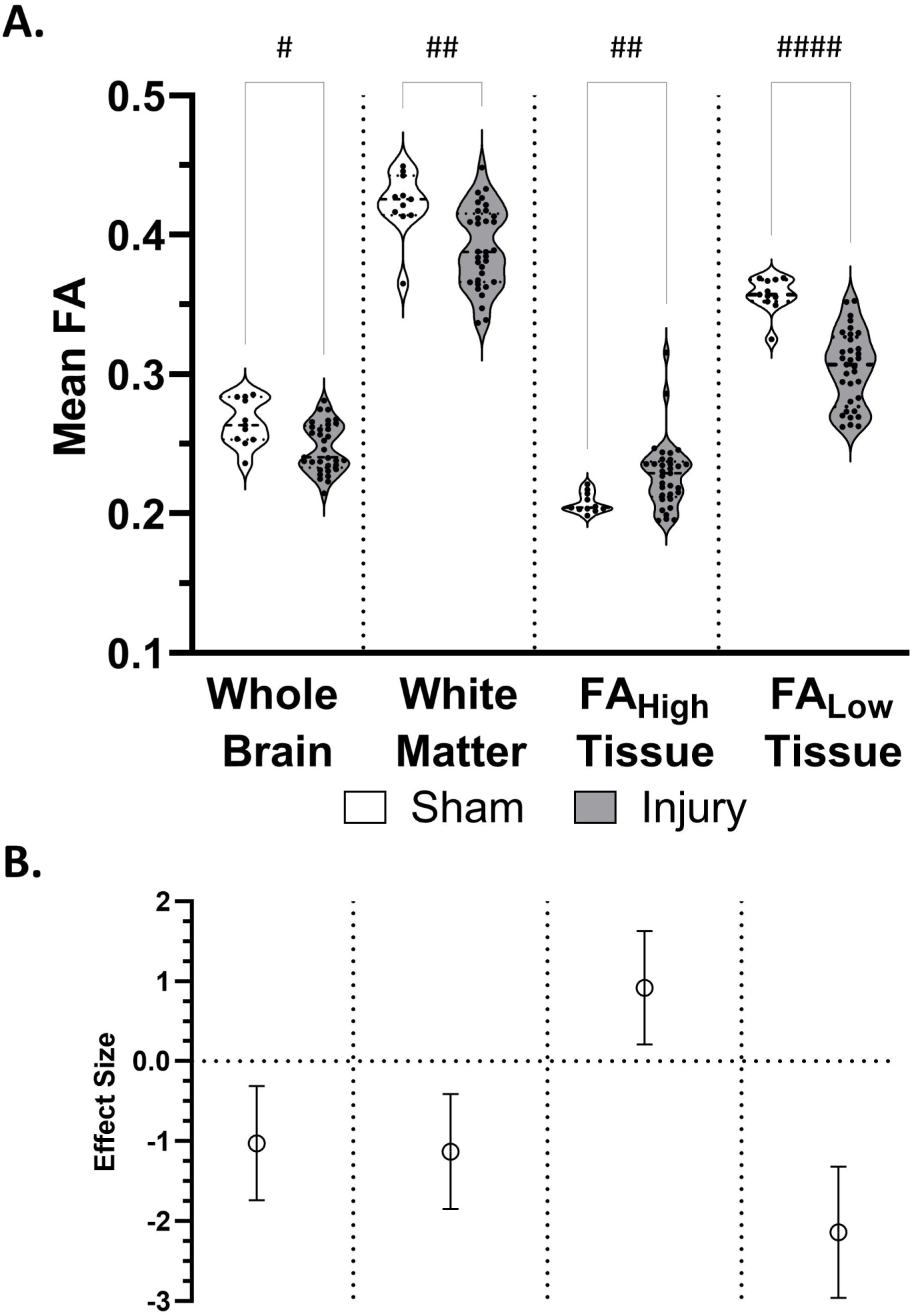
Data-driven versus whole brain ROI-derived mean FA measurements at 1m post-injury. Mean FA of whole brain segmented by tissue type for individual rats for [**A**] Whole Brain (grey and white matter), White Matter (whole brain white matter only), and Pathologic Tissue (a data-derived region of altered FA voxels based on any voxel being significantly lower or higher than the corresponding sham voxels in at least 10% of the rats measured in the group. [**B**] Graph of effect size (Hedges’ g) with confidence intervals for the given comparisons. Mann-Whitney U, #-p<0.05, ##-p<0.01, ####-p<0.0001.

We next tested the same 1mo data for group effects that were collected using a series of 7, pre-selected, bilateral, small ROIs that were located within brain regions expected to be affected by the injury, based either on distance from, or connectivity to the primary impact site. Using non-parametric statistics due to the non-normally distributed data, no ROI group comparison survived correction for multiple ROI comparisons (n=14 bilateral ROIs), despite a trend toward lower FA throughout all regions in injured compared to the sham group at 1mo post-injury (**Fig. 4A-D, I-L**). Effect sizes for the group comparison varied largely by distance from the primary impact zone, for example it peaked at -1.43 in the left auditory cortex grey matter that is adjacent to the primary impact, -1.02 in the left hippocampus immediately ventral to the major impact zone, and -1.05 in the adjacent genu of the corpus callosum, but it ranged from -0.30 to -0.91 in more distant, ipsilateral posterior or contralateral regions (**Fig. 4E-L**). We found similar or smaller effect sizes for the other tensor scalars: mean, axial and radial diffusivity (S7-9). Therefore, while the effect size of the primary impact site ROI approached the data driven comparison for FA_LOW_ (-1.43 versus 2.2), all other ROIs were smaller.

**Fig. 4.**
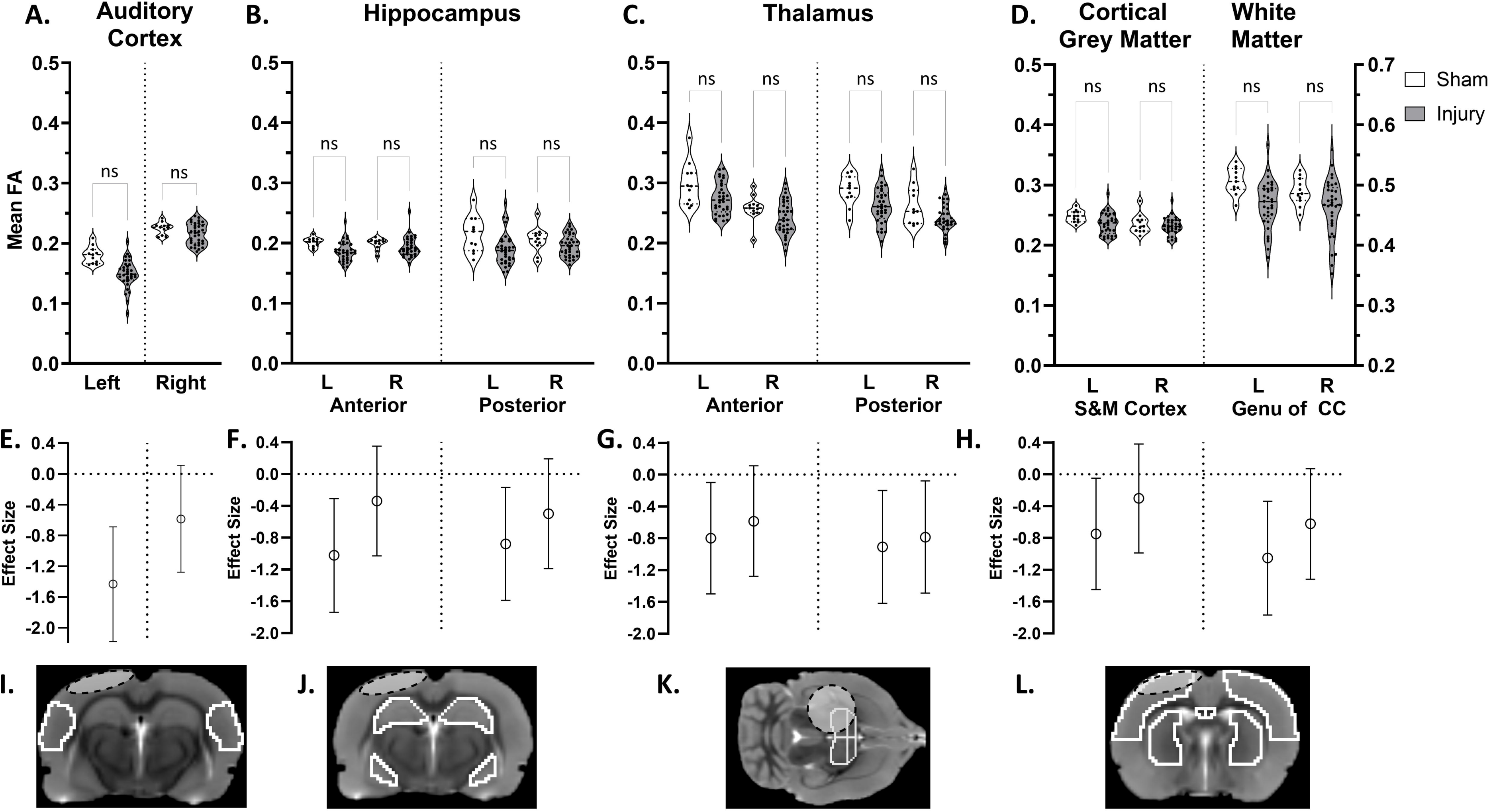
ROI Brain Parcellation Analysis of Mean FA at 1month post-injury. Mean FA in sham and injured rats (open and closed bars respectively) in: [**A**] the auditory cortex, the site of the primary injury, [**B**] anterior and poster hippocampus, [**C**] anterior and posterior thalamus, [**D**] sensorimotor (S&M) cortex and genu of the corpus callosum (CC). [**E-H**] Graphs of group effect size with confidence interval for the ROIs. [**I-L**] Locations of the ROIs. White plots are sham group, and grey plots are injury group. Grey disk represents the location of craniectomy. Key: L- left, R- right, NS-Not significant. None of the trends of lower FA in the injury survived correction for multiple corrections (Kruskal-Wallis test, Dunn’s multi-ROI correction).

### Post-Injury, Longitudinal Comparison of Methods

We took the 3 ROIs with the highest effect size when compared to shams, (ROIs with effect size >1 for the FA analysis, **Figs. 3 and 4**), the top 3 ROIs with the largest effect size for the other tensor scalars, and the whole brain and whole brain white matter ROIs (**S10-12**) and compared their ability to characterize the longitudinal changes in tensor scalars over time post-injury (2d. to 5m., **Fig. 5**) to data obtained from the data-driven method. The latter data were obtained by comparison to mean and standard deviation sham population data calculated over all post-injury time-points in order to incorporate any age-related changes in diffusion scalars. A mixed-effect ANOVA model showed no significant, overall time effect for any of the ROI data comparisons (**Fig. 5A-E**), but it was significant for the data-derived, pathologic tissue composed of FA_High_ and FA_Low_ values (F(2.321, 72.74)=10.64, P<.0001; F(2.014, 63.11)=15.61, P<.0001, for FA_High_ and FA_Low_ respectively). Here there was a significant change from 2-9d. and 2d-1m., and 1-5m. in FA_High_ tissue, and 2d-1m., 9d-1m., and 1-5m. in FA_Low_ tissue, post-injury when adjusted for multiple comparisons of time-points (p<.05, **Fig. 5F-G**). The other tensor scalars showed quite different effects compared to FA, with temporal significance reached among all ROIs analyzed, especially in the injury adjacent auditory cortex (**S10-12**). However, the data-driven analysis also showed significance over time and in the same direction as the ROIs.

**Fig. 5.**
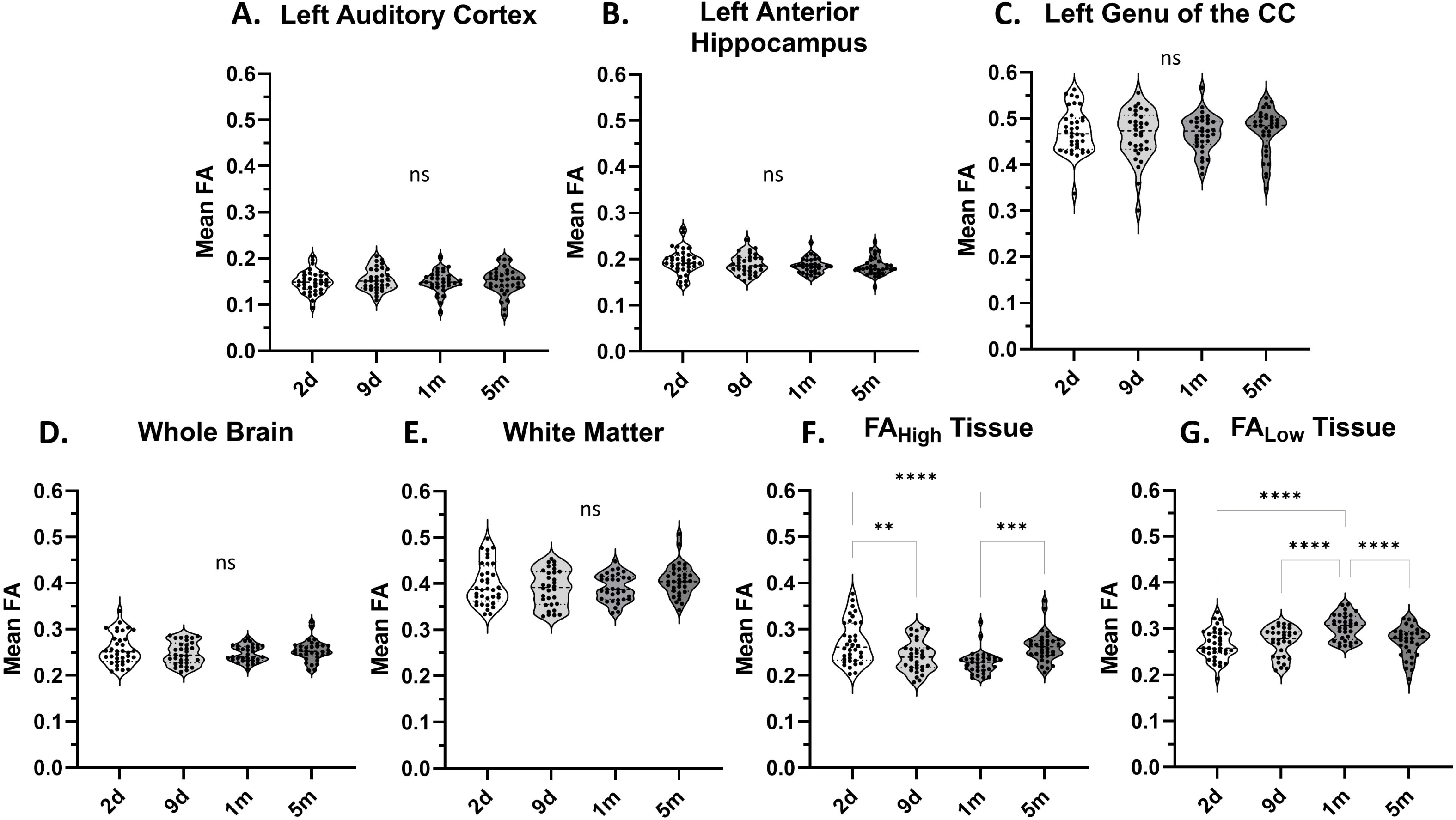
Comparison of data-driven vs ROI-derived changes in FA longitudinally after injury. Mean regional FA changes from 2d. to 5mo. after LFPI for [**A-F**] ROIs with an effect size greater than 1 at 1m post-LFPI (see Figs. 3 and 4) compared to [**F**] values derived using a data-driven method. Key: **,***,**** = Mixed-effects Tukey corrected multiple comparisons, p<0.01, p<0.001, p<0.0001; ns = no significant comparisons.

We computed overlap maps of the data-driven FA-derived volumes at each post-injury time-point to determine the location of the pathologic tissue volumes identified by FA_High_ and FA_Low_ values (**Fig. 6**). Apart from the initial 2d post-injury time-point, the data showed marked similarity between the locations of both FA volumes across each time-point and a relatively high degree of overlap between injured rats (**Fig. 6A**). Heavily overlapping FA_Low_ regions were constrained almost exclusively to the ipsilateral corpus callosum, and medial and superficial grey matter cortical areas, while highly overlapping FA_High_ regions occurred chiefly within ipsilateral, deep cortical grey matter. Notable temporal changes were largely related to regions of FA_Low_ which more chronically developed larger areas of overlap between rats within the contralateral corpus callosum, dorsal hippocampus, and in the ipsilateral cerebellum indicative of continued white matter atrophy (**Fig. 6B**). There were also marked temporal volume changes for the other scalars, and similar to FA, they all showed relatively tight confidence limits compared to sham, indicating the robustness of the methodology (**S13-15**).

**Fig. 6.**
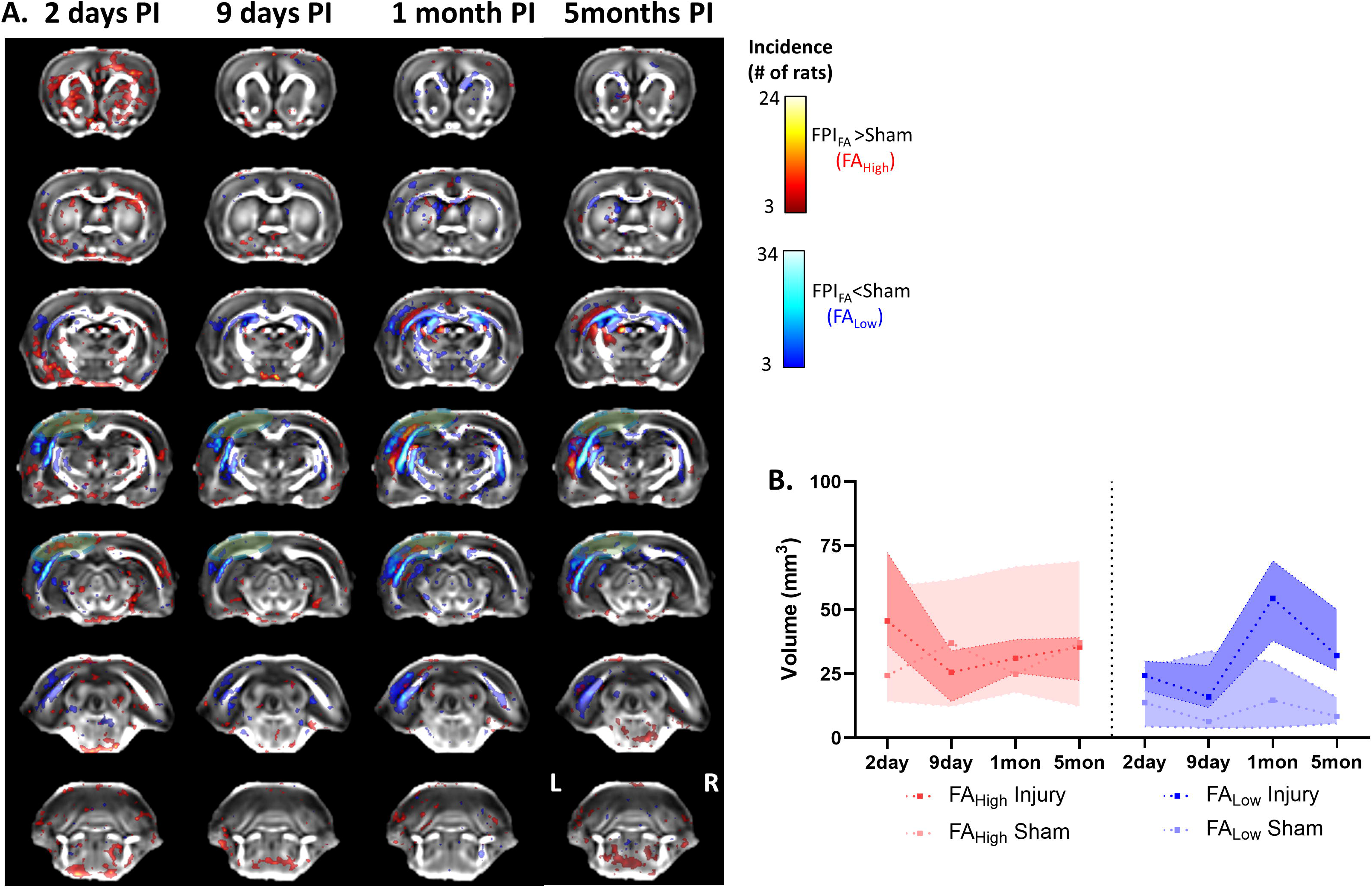
Data-driven FA-derived volumetric changes over time. **[A]** Population overlay maps for volumes of FA_High_ and FA_Low_ tissue. Each column shows the change in pathologic tissue in the injured group across time, at 2d., 9d., 1mo. and 5mos. post injury, overlayed on mean FA map. **<u>Key:</u>** L- Left, R- Right, Green disk represents the location of craniectomy. **[B]** Summary of volumes of pathologic tissue identified by FA_High_ or FA_Low_ across time in both injury and sham animals. Data are medians with 95% confidence intervals.

Visual comparison of the temporal changes for all low and high volumes of each tensor scalars revealed quite different trajectories from each, with mean, radial and axial diffusivity volumes being the most similar, and FA showing very different changes (**Fig. 7**). The volume of pathologic tissue that was significantly higher in the injured animals when compared to shams varied across all post injury time-points. FA_High_ tissue volume was never significantly different in injured rats compared to shams, whereas the volume of FA_Low_ tissue became significantly higher in injured animals at 1mo and remained higher at 5m post injury. For the other diffusion scalars (MD, AD, and RD) the volume of Scalar_High_ tissue became significantly larger in the injured animals at 1mo. post injury and remained high up to 5m. While the volumes of Scalar_Low_ tissue were initially significantly higher in injured animals at 2ds. post injury, there were no differences from 9d. onwards after injury (**Table 1**, **Fig. 7**). There was a significant overall time effect for all scalar volumes (**Table 2**, P<0.05). The greatest change in all Scalar_High_ volumes occurred between 2d and 1m (Effect size -1.35 to 1.60, P<0.0001) while for Scalar_Low_ volumes it occurred earlier, between 2-9d (Effect size -0.98 to 1.36, P<0.001). The greatest changes in FA_High_ occurred 2d-9d (Effect size -0.655, P<0.05 where FA_Low_ occurred later at 2d-1m (Effect Size 1.35, P<0.0001).

**Fig. 7.**
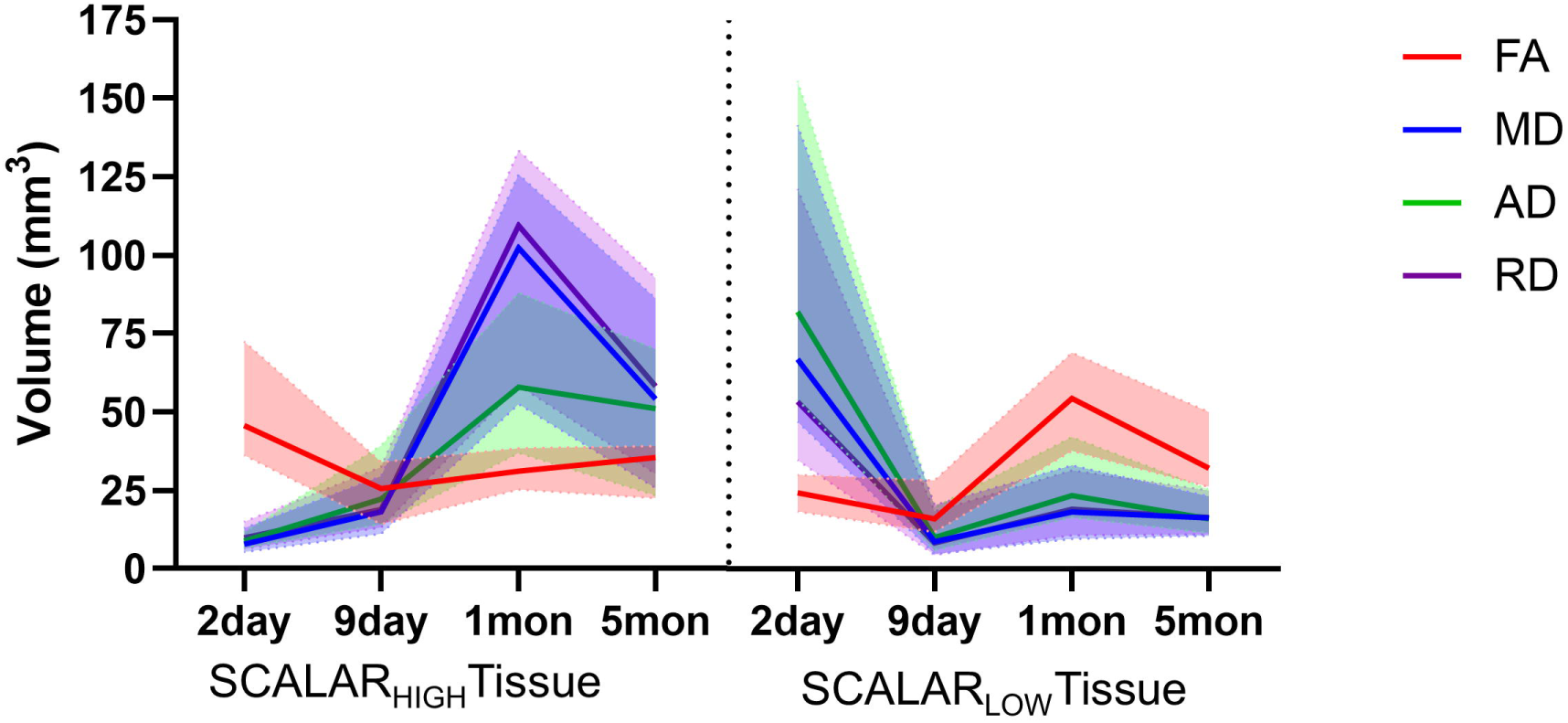
Summary of volumes of diffusion metric altered tissue changes over time. The changes in volumes of tissue with significantly higher or lower than values of voxel-wise derived diffusion scalar metrics: fractional anisotropy (FA), mean diffusion (MD, axial diffusion (AD), and radial diffusion (RD), as compared to sham animals.

**Table 1.**
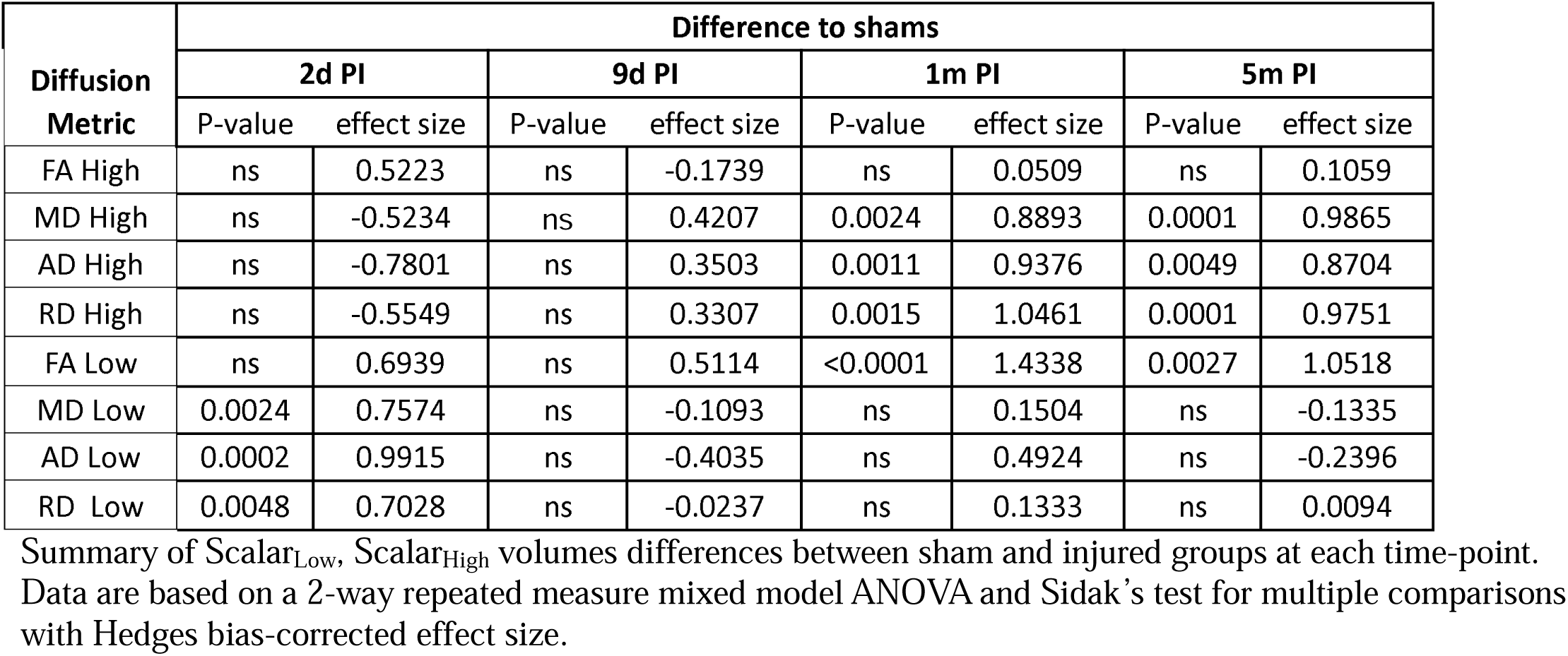
Difference between shams and TBI rats in volume of pathologic tissue at 2d, 9d, 1m, and 5m post injury.

| Diffusion<br>Metric | Difference to shams |  |  |  |  |  |  |  |
| --- | --- | --- | --- | --- | --- | --- | --- | --- |
|  | 2d PI |  | 9d PI |  | 1m PI |  | 5m PI |  |
|  | P-value | effect size | P-value | effect size | P-value | effect size | P-value | effect size |
| FA High | ns | 0.5223 | ns | -0.1739 | ns | 0.0509 | ns | 0.1059 |
| MD High | ns | -0.5234 | ns | 0.4207 | 0.0024 | 0.8893 | 0.0001 | 0.9865 |
| AD High | ns | -0.7801 | ns | 0.3503 | 0.0011 | 0.9376 | 0.0049 | 0.8704 |
| RD High | ns | -0.5549 | ns | 0.3307 | 0.0015 | 1.0461 | 0.0001 | 0.9751 |
| FA Low | ns | 0.6939 | ns | 0.5114 | <0.0001 | 1.4338 | 0.0027 | 1.0518 |
| MD Low | 0.0024 | 0.7574 | ns | -0.1093 | ns | 0.1504 | ns | -0.1335 |
| AD Low | 0.0002 | 0.9915 | ns | -0.4035 | ns | 0.4924 | ns | -0.2396 |
| RD Low | 0.0048 | 0.7028 | ns | -0.0237 | ns | 0.1333 | ns | 0.0094 |

Summary of $\text{Scalar}_{\text{Low}}$ , $\text{Scalar}_{\text{High}}$ volumes differences between sham and injured groups at each time-point. Data are based on a 2-way repeated measure mixed model ANOVA and Sidak's test for multiple comparisons with Hedges bias-corrected effect size.

**Table 2.**
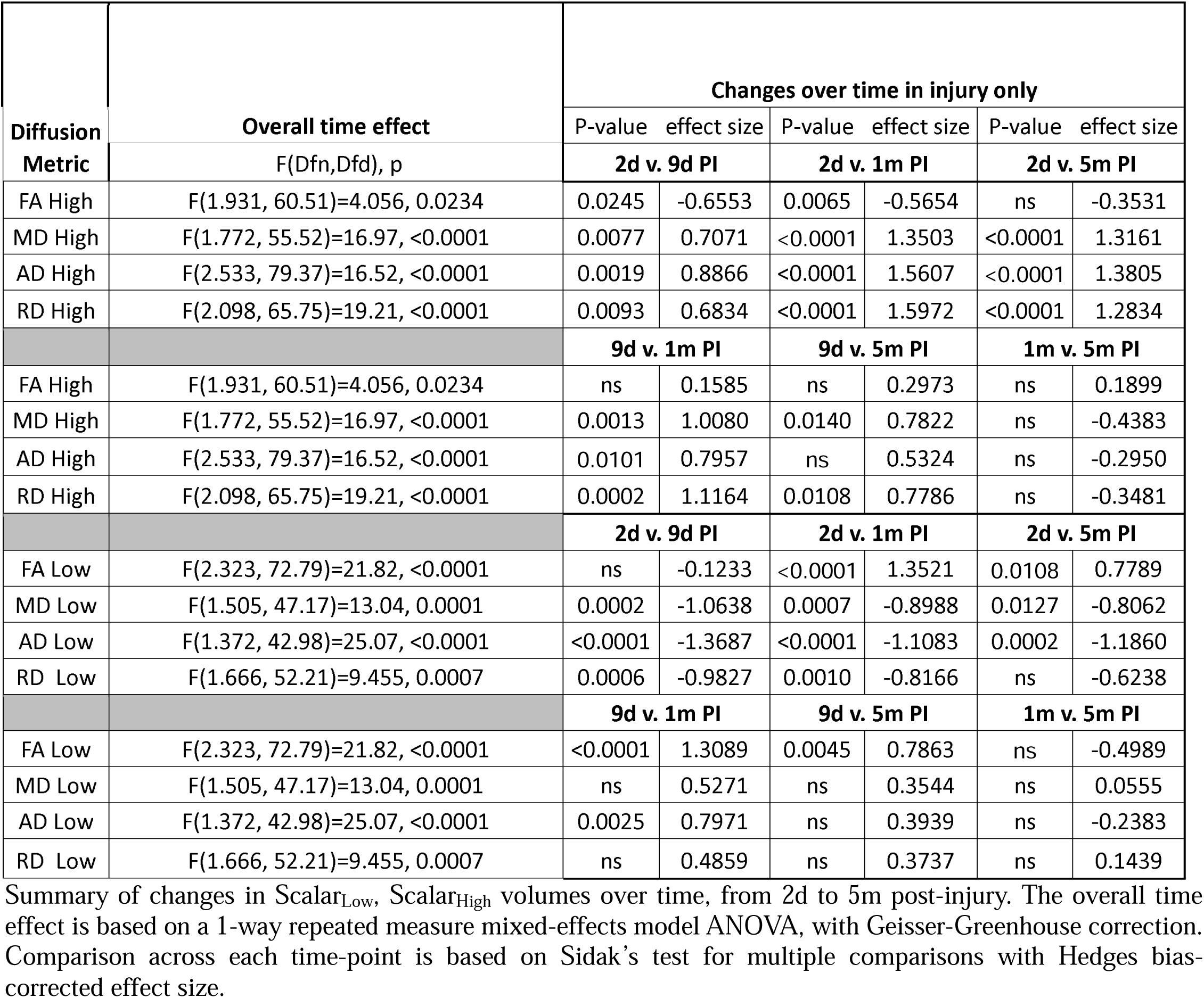
Changes overtime in volume of pathologic tissue in injured animals.

## Discussion

The results show that this data-driven, unbiased z-score-based approach successfully delineated volumes of pathologic grey and white matter in the preclinical injured brain as determined by volumes of interest delineated by FA_HIGH_ and FA_LOW_. This effect was also consistent across the other tensor scalars and overall resulted in excellent effect sizes at >80% power that allowed discrimination between injured and sham control groups. The approach was also shown to be consistent and robust across multiple time-points post-LFPI with both FA and with all tensor-derived scalars. The group effect size of the scalar values underlying each data-driven volume of interest was at, or close to the greatest effect size of a ROI-based approach using brain areas typically affected by injury, indicating the robustness of the effect.

An ROI-based approach to image analysis using a priori information to site ROIs normally affords the greatest sensitivity to detect differences to sham or naïve control subjects. However, this presupposes that all information is known, is accurate, and that no other brain regions are important for delineating any effect of injury or intervention. This approach may lead to missed opportunities for finding pathology that is unexpected, and that may otherwise be important for ascribing altered structure or function that could useful for interpreting other data acquired from the same subjects. In addition, using typically located ROIs may not be as sensitive to a potential treatment effect because limiting the effect of injury to regions close to the primary impact site that normally provide a large effect size, may washout any smaller changes amounting to a therapeutic effect. The data-driven, voxel-based approach is agnostic to prior information and enables the derivation of regions of pathology unbiased by subjective information. This provides a more flexible approach to identify unexpected regions of pathology while maintaining an effect size approaching the level of the most sensitive ROIs for detecting the effect of injury. The method can easily be applied to most, if not all, parametric maps of MRI metrics describing pathology after injury or other overt pathology that is not focal. Furthermore, unlike a ROI approach where the heterogeneity of injury may washout any effect, the data-driven method localizes areas of pathology by interrogating the whole brain so that there is no requirement for overlapping areas of pathology across subjects in order to obtain data sensitive to the effect of injury.

Recent clinical findings that have used this data-driven technique indicate the importance of applying the correction factor to the data(Gimbel et al., 2024b, 2024a; Mayer et al., 2024). This study has shown that the technique can be readily extended to preclinical studies to systematically identify regions of pathology over time post-injury. The distillation of pathology into two univariate data volumes per brain (Scalar_LOW_, Scalar_HIGH_) captures the extent of injury affecting the whole brain. This is particularly relevant for comparison of imaging findings to plasma or serum biomarker analysis in the same subject. The washout of brain-specific proteins to the systemic circulation is unlikely to be limited to small brain regions reported on by discrete ROIs. A much larger part of the brain is likely to be involved, as indicated by the widespread opening of the blood brain-barrier after FPI injury(Lin et al., 2012) and in multiple other models(Li et al., 2017; Rubovitch et al., 2011; Schneider et al., 2002; Zvenigorodsky et al., 2024). As such, the analysis method detailed herein that produces an outcome of whole brain burden of injury, rather than a more limited ROI analysis, is best suited for studies seeking to validate novel blood markers for predicting injury severity or outcome by using image analysis as ground truth. Other potential practical implications for future preclinical TBI studies for this whole brain analysis would be a role in evaluating therapeutic interventions using a single number (eg. Scalar_LOW_ volume). In addition, whole brain volumes of pathology could be useful predictive biomarkers for behavioral outcome, especially when the volume is limited to specific circuits or structural locations known to underpin behavioral deficits.

### Requirements, Limitations and Potential Solutions

Similar to most image post-processing pipelines, there is a critical requirement to conduct precise image co-registration to enable accurate computation of sham group mean and standard deviation data for computation of z-score maps. This also requires that all injury data be consistently aligned with the study template. However, when the injury is very severe, false positive, significant z-scores can occur in misaligned voxels. This can be mitigated to some extent by incorporating masks to exclude severe pathology from the calculation of the warp and affine transformations for coregistration. However, the majority of image coregistration algorithms use the spatial information contained within the greyscale intensity so that errors can occur when there are large bright areas within an injured brain, for example, edema or cavity on T2-weighted image data. We chose to use FA data for the coregistration due to the very distinct differences in grey and white matter intensities that aid the accuracy of the registration. However, injured grey matter at early times after injury may become hyperintense indicated by high FA, which can erroneously be coregistered to white matter within a mean template image. While we manually checked all data and corrected any problems in alignment quality, it does make the analysis process longer. One possibility to resolve or at least help mitigate this is to use coregistration algorithms that leverage the higher order information contained within the calculated diffusion tensor images to improve the accuracy of the registration (Zhang et al., 2006). Another possibility, with the downside of increased scanning time, is to use higher resolution scan acquisition parameters to capture more detailed microstructural information. This can then be leveraged by the co-registration routines to improve image alignment and enhance experimental rigor.

Other potential issues are spatially incomplete image data i.e. partial brains, and the presence of image artifacts due to hardware or acquisition problems. This unbiased analysis used herein requires equal representation of all brain voxels from across all subjects. This is practically taken care of by generating a common brain mask over the entire sample population to ensure that all computed z-statistics are created from voxels representing the same number of subjects. If one spatially incomplete brain is included, then the whole group analysis is reduced to the region of partial brain that is common to all subjects within the study population, severely limiting the analysis. This can be avoided by careful attention to input data through image acquisition of the same regions of brain during scanning by reference to brain and extracranial landmarks. The presence of image artifacts has a similar effect; the occurrence of image ghosting due to aliasing from Nyquist ghosting or chemical shift artifacts can result in inflated z-scores. These can be mitigated by including only the highest quality data, or if the problems are in sham data, excluding those data where the artifact occurs in regions anticipated to be involved in injury. However, we generally avoided excluding data unless the scan acquisition completely failed, since the presence of artifacts are likely to occur equally across all subjects, regardless of group membership. While this promotes more noise in the final analysis and contributes to the labelling of artifacts as injury in both injured and shams groups, it is consistent with the unbiased nature of the technique. The technique has recently been shown to be easily applicable to the controlled cortical impact (CCI) model, and across multiple sites using data harmonization techniques (Kislik et al., 2025).

### Interpretability of Longitudinal Volumes of Pathology

The longitudinal data analysis of volumes of microstructure pathology described by MD_LOW_, AD_LOW_ and RD_LOW_ were highest early after LFPI injury, but thereafter remained relatively unchanged compared to shams. Meanwhile there was a corresponding increase in these Scalar_HIGH_ volumes above sham values at later time points, consistent with a typical pseudo-normalization of diffusion parameters that was first reported in the stroke literature(Lansberg et al., 2001). This temporal transformation is in agreement with chronic resolution of edema, tissue atrophy and cavity formation in this model(Obenaus et al., 2007; Smith et al., 2023). Large regions of MD_LOW_ early after LFPI have been measured before and are interpretable as cytotoxic edema and relate to metabolic disruption(Smith et al., 2023). The highest volume of RD_HIGH_ occurred at 1month post-injury consistent with the known time-course of demyelination in this and other models (Armstrong et al., 2016; Chary et al., 2021). The volume of FA_LOW_ was consistently above shams from the first post-injury measurement, and thereafter the trajectory increased consistent with on-going axonal degeneration that was highest at 1month but remained high at 5 months. Indeed, this agrees with the finding of reduced apparent fiber density, track density and average pathlengths in corpus collosum at 3 months post-FPI injury(Wright et al., 2017), as well as reduced FA and increased MD, AD and RD at the same timepoint. The interpretation of the early increase in FA_HIGH_ volumes is not straightforward since it indicates increased alignment of diffusion trajectories in a field gradient which, at first glance might not be explainable on a microstructural level. This finding also been observed in the CCI model(Harris et al., 2016). Given that 90% white matter voxels contain crossing fibers (Jeurissen et al., 2013), deafferentation could be expected to give rise to high FA. However, a readout of this – the tensor mode shape analysis was not associated with regions of FA_HIGH_ after CCI, and in the current data, FA_HIGH_ was constrained to cortical grey matter and subcortical structures. Regions of increased FA were found to be consistent with the coherent organization of reactive astrocytes (Budde et al., 2011). It will be important in future work to determine the relationship of immunohistology-level microstructural changes to these derived volumes of pathology. To that end, there are ongoing efforts by the Translational Outcomes Project in Neurotrauma Consortium (TOPNT) to characterize the detailed pathology underpinning the regions identified by this data driven technique in the CCI and FPI rat models using image aligned histological assessment (Radabaugh et al., 2025; Wanner et al., 2025). Regardless of the exact biological interpretation of these proxy markers of pathology, the presence of different trajectories of each scalar volume post-injury confirms the idea that they represent separable, pathologic processes that can be tracked temporally.

Given the current use of this image processing technique or techniques like it across the clinical TBI field, the interest by funding bodies to translate biomarkers across species wherever possible, and the statistical robustness and reproducibility for its use within an experimental model of TBI, we would recommend that the preclinical TBI field might include it within existing imaging pipelines for correlation to, and validation of accompanying biomarkers such as biofluids and behavioral readouts.

To conclude, we have shown that this relatively simple data-driven technique, which is as sensitive as a ROI analysis for detecting group differences, can be easily deployed for preclinical data analysis to generate unbiased measures of diffusion scalar metrics through an unbiased volume analysis of the whole brain.

## Acknowledgments

We would like to thank Dr Andrew Frew and the UCLA In Vivo MR Imaging and Spectroscopy Center.

## Authorship Confirmation/ Contribution

**Gregory Smith:** Data curation, Formal Analysis, Writing-Original draft preparation, Software, Visualization. **Cesar Santana-Gomez.**: Data curation, Writing-Reviewing and Editing. **Richard Staba:** Writing-Reviewing and Editing, Funding acquisition, Project administration, Resources. **Neil Harris:** Conceptualization, Methodology, Formal Analysis, Writing-Original draft preparation, Supervision, Funding acquisition, Resources.

## Figure Legends

**Supplemental Fig. 1. Data-driven Analysis of MD at 1month after injury.** [**A]** Volumes of MD_High_ and MD_Low_ in sham and LFPI rats. [**B]** A population incidence map of MD_High_ and MD_Low_ volumes from all injured rats highlighting the location of altered tissue (voxel pseudocolors indicate number of rats), overlayed on mean FA map. [**C]** Graph of statistical effect size, (Hedges’ g) with confidence intervals for the effect of group for MD_High_ and MD_Low_ volumes. Kruskal-Wallis test ### p<0.001. Green disk represents the location of craniectomy.

**Supplemental Fig. 2. Data-driven Analysis of AD at 1month after injury.** [**A]** Volumes of AD_High_ and AD_Low_ in sham and LFPI rats. [**B]** A population incidence map of AD_High_ and AD_Low_ volumes from all injured rats highlighting the location of altered tissue (voxel pseudocolors indicate number of rats), overlayed on mean FA map. [**C]** Graph of statistical effect size, (Hedges’ g) with confidence intervals for the effect of group for AD_High_ and AD_Low_ volumes. Kruskal-Wallis test ## p<0.01. Green disk represents the location of craniectomy.

**Supplemental Fig. 3. Data-driven Analysis of RD at 1month after injury.** [**A]** Volumes of RD_High_ and RD_Low_ in sham and LFPI rats. [**B]** A population incidence map of RD_High_ and RD_Low_ volumes from all injured rats highlighting the location of altered tissue (voxel pseudocolors indicate number of rats), overlayed on mean FA map. [**C]** Graph of statistical effect size, (Hedges’ g) with confidence intervals for the effect of group for RD_High_ and RD_Low_ volumes. Kruskal-Wallis test ### p<0.001. Green disk represents the location of craniectomy.

**Supplemental Fig. 4. Data-driven versus whole brain ROI-derived mean MD measurements at 1m post-injury.** Mean MD of whole brain segmented by tissue type for individual rats for [**A**] Whole Brain (grey and white matter), White Matter (whole brain white matter only), and Pathologic Tissue (a data-derived region of altered MD voxels based on any voxel being significantly lower or higher than the corresponding sham voxels in at least 10% of the rats measured in the group. [**B**] Graph of effect size (Hedges’ g) with confidence intervals for the given comparisons. Mann-Whitney U, ###-p<0.001.

**Supplemental Fig. 5. Data-driven versus whole brain ROI-derived mean AD measurements at 1m post-injury.** Mean AD of whole brain segmented by tissue type for individual rats for [**A**] Whole Brain (grey and white matter), White Matter (whole brain white matter only), and Pathologic Tissue (a data-derived region of altered AD voxels based on any voxel being significantly lower or higher than the corresponding sham voxels in at least 10% of the rats measured in the group. [**B**] Graph of effect size (Hedges’ g) with confidence intervals for the given comparisons. Mann-Whitney U, ##-p<0.01, ###-p<0.001.

**Supplemental Fig. 6. Data-driven versus whole brain ROI-derived mean RD measurements at 1m post-injury.** Mean RD of whole brain segmented by tissue type for individual rats for [**A**] Whole Brain (grey and white matter), White Matter (whole brain white matter only), and Pathologic Tissue (a data-derived region of altered RD voxels based on any voxel being significantly lower or higher than the corresponding sham voxels in at least 10% of the rats measured in the group. [**B**] Graph of effect size (Hedges’ g) with confidence intervals for the given comparisons. Mann-Whitney U, #-p<0.05, ####-p<0.0001.

**Supplemental Fig. 7. ROI Brain Parcellation Analysis of Mean MD at 1month post-injury.** Mean MD in sham and injured rats (open and closed bars respectively) in: [**A**] the auditory cortex, the site of the primary injury, [**B**] anterior and poster hippocampus, [**C**] anterior and posterior thalamus, [**D**] sensorimotor (S&M) cortex and genu of the corpus callosum (CC). [**E-H**] Graphs of group effect size with confidence interval for the ROIs. [**I-L**] Locations of the ROIs. White plots are sham group, and grey plots are injury group. Grey disk represents the location of craniectomy. Key: L-left, R-right, NS-Not significant. None of the trends of altered MD in the injury survived correction for multiple corrections (Kruskal-Wallis test, Dunn’s multi-ROI correction).

**Supplemental Fig. 8. ROI Brain Parcellation Analysis of Mean AD at 1month post-injury.** Mean AD in sham and injured rats (open and closed bars respectively) in: [**A**] the auditory cortex, the site of the primary injury, [**B**] anterior and poster hippocampus, [**C**] anterior and posterior thalamus, [**D**] sensorimotor (S&M) cortex and genu of the corpus callosum (CC). [**E-H**] Graphs of group effect size with confidence interval for the ROIs. [**I-L**] Locations of the ROIs. White plots are sham group, and grey plots are injury group. Grey disk represents the location of craniectomy. Key: L-left, R-right, NS-Not significant. None of the trends of altered AD in the injury survived correction for multiple corrections (Kruskal-Wallis test, Dunn’s multi-ROI correction).

**Supplemental Fig. 9. ROI Brain Parcellation Analysis of Mean RD at 1month post-injury.** Mean RD in sham and injured rats (open and closed bars respectively) in: [**A**] the auditory cortex, the site of the primary injury, [**B**] anterior and poster hippocampus, [**C**] anterior and posterior thalamus, [**D**] sensorimotor (S&M) cortex and genu of the corpus callosum (CC). [**E-H**] Graphs of group effect size with confidence interval for the ROIs. [**I-L**] Locations of the ROIs. White plots are sham group, and grey plots are injury group. Grey disk represents the location of craniectomy. Key: L-left, R-right, NS-Not significant. None of the trends of altered RD in the injury survived correction for multiple corrections (Kruskal-Wallis test, Dunn’s multi-ROI correction).

**Supplemental Fig. 10. Comparison of data-driven vs ROI-derived changes in MD longitudinally after injury.** Mean regional MD changes from 2d. to 5mo. after LFPI for [**A-E**] Top three ROIs based on effect size as compared to shams at 1m post-LFPI (see Sup. Figs. 2 and 3) and whole brain, and white matter only as compared to [**F-G**] values derived using a data-driven method. Key: *,**,***,**** = Mixed-effects Tukey corrected multiple comparisons, p<0.05, p<0.01,p<0.001, p<0.0001; ns = no significant comparisons.

**Supplemental Fig. 11. Comparison of data-driven vs ROI-derived changes in AD longitudinally after injury.** Mean regional AD changes from 2d. to 5mo. after LFPI for [**A-E**] Top three ROIs based on effect size as compared to shams at 1m post-LFPI (see Sup. Figs. 2 and 3) and whole brain, and white matter only as compared to [**F-G**] values derived using a data-driven method. Key: *,**,***,**** = Mixed-effects Tukey corrected multiple comparisons, p<0.05, p<0.01,p<0.001, p<0.0001; ns = no significant comparisons.

**Supplemental Fig. 12. Comparison of data-driven vs ROI-derived changes in RD longitudinally after injury.** Mean regional RD changes from 2d. to 5mo. after LFPI for [**A-E**] Top three ROIs based on effect size as compared to shams at 1m post-LFPI (see Sup. Figs. 2 and 3) and whole brain, and white matter only as compared to [**F-G**] values derived using a data-driven method. Key: *,**,***,**** = Mixed-effects Tukey corrected multiple comparisons, p<0.05, p<0.01,p<0.001, p<0.0001; ns = no significant comparisons.

**Supplemental Fig. 13. Data-driven MD-derived volumetric changes over time. [A]** Population overlay maps for volumes of MD_High_ and MD_Low_ tissue. Each column shows the change in pathologic tissue in the injured group across time, at 2d., 9d., 1mo. and 5mos. post injury, overlayed on mean FA map. **<u>Key:</u>** L- Left, R- Right, Green disk represents the location of craniectomy. **[B]** Summary of volumes of pathologic tissue identified by MD_High_ or MD_Low_ across time in both injury and sham animals. Data are medians with 95% confidence intervals.

**Supplemental Fig. 14. Data-driven AD-derived volumetric changes over time. [A]** Population overlay maps for volumes of AD_High_ and AD_Low_ tissue. Each column shows the change in pathologic tissue in the injured group across time, at 2d., 9d., 1mo. and 5mos. post injury, overlayed on mean FA map. **<u>Key:</u>**L- Left, R- Right, Green disk represents the location of craniectomy. **[B]** Summary of volumes of pathologic tissue identified by AD_High_ or AD_Low_ across time in both injury and sham animals. Data are medians with 95% confidence intervals.

**Supplemental Fig. 15. Data-driven RD-derived volumetric changes over time. [A]** Population overlay maps for volumes of RD_High_ and RD_Low_ tissue. Each column shows the change in pathologic tissue in the injured group across time, at 2d., 9d., 1mo. and 5mos. post injury, overlayed on mean FA map. **<u>Key:</u>** L- Left, R- Right, Green disk represents the location of craniectomy. **[B]** Summary of volumes of pathologic tissue identified by RD_High_ or RD_Low_ across time in both injury and sham animals. Data are medians with 95% confidence intervals.

## References

Amadi, D., Kiwuwa-Muyingo, S., Bhattacharjee, T., Taylor, A., Kiragga, A., Ochola, M., Kanjala, C., Gregory, A., Tomlin, K., Todd, J., Greenfield, J., 2024. Making Metadata Machine-Readable as the First Step to Providing Findable, Accessible, Interoperable, and Reusable Population Health Data: Framework Development and Implementation Study. Online J Public Health Inform 16, e56237. 10.2196/56237

Andersson, J.L.R., Graham, M.S., Zsoldos, E., Sotiropoulos, S.N., 2016. Incorporating outlier detection and replacement into a non-parametric framework for movement and distortion correction of diffusion MR images. NeuroImage 141, 556–572. 10.1016/j.neuroimage.2016.06.058

Andersson, J.L.R., Skare, S., Ashburner, J., 2003. How to correct susceptibility distortions in spin-echo echo-planar images: application to diffusion tensor imaging. Neuroimage 20, 870–888. 10.1016/S1053-8119(03)00336-7

Andersson, J.L.R., Sotiropoulos, S.N., 2016. An integrated approach to correction for off-resonance effects and subject movement in diffusion MR imaging. NeuroImage 125, 1063–1078. 10.1016/j.neuroimage.2015.10.019

Armstrong, R.C., Mierzwa, A.J., Marion, C.M., Sullivan, G.M., 2016. White matter involvement after TBII]: Clues to axon and myelin repair capacity. Experimental Neurology 275, 328–333. 10.1016/j.expneurol.2015.02.011

Avants, B.B., Epstein, C.L., Grossman, M., Gee, J.C., 2008. Symmetric diffeomorphic image registration with cross-correlation: evaluating automated labeling of elderly and neurodegenerative brain. Med Image Anal 12, 26–41. 10.1016/j.media.2007.06.004

Begley, C.G., Ioannidis, J.P.A., 2015. Reproducibility in Science. Circulation Research 116, 116–126. 10.1161/CIRCRESAHA.114.303819

Budde, M.D., Janes, L., Gold, E., Turtzo, L.C., Frank, J.A., 2011. The contribution of gliosis to diffusion tensor anisotropy and tractography following traumatic brain injuryI]: validation in the rat using Fourier analysis of stained tissue sections 2248–2260. 10.1093/brain/awr161

Chary, K., Nissi, M.J., Nykänen, O., Manninen, E., Rey, R.I., Shmueli, K., Sierra, A., Gröhn, O., 2021. Quantitative susceptibility mapping of the rat brain after traumatic brain injury. NMR Biomed 34, e4438. 10.1002/nbm.4438

Collins, F.S., Tabak, L.A., 2014. Policy: NIH plans to enhance reproducibility. Nature 505, 612–613. 10.1038/505612a

Cordero-Grande, L., Christiaens, D., Hutter, J., Price, A.N., Hajnal, J.V., 2019. Complex diffusion-weighted image estimation via matrix recovery under general noise models. NeuroImage 200, 391–404. 10.1016/j.neuroimage.2019.06.039

Davenport, N.D., Lim, K.O., Armstrong, M.T., Sponheim, S.R., 2012. Diffuse and spatially variable white matter disruptions are associated with blast-related mild traumatic brain injury. Neuroimage 59, 2017–2024. 10.1016/j.neuroimage.2011.10.050

Fox, R., Santana-Gomez, C., Shamas, M., Pavade, A., Staba, R., Harris, N.G., 2024. Different Trajectories of Functional Connectivity Captured with Gamma-Event Coupling and Broadband Measures of Electroencephalographic in the Rat Fluid Percussion Injury Model. Neurotrauma Reports 5, 1027–1038. 10.1089/neur.2024.0045

Gimbel, S.I., Hungerford, L., Twamley, E.W., Ettenhofer, M.L., 2024a. Authors’ Response to “Revisiting Subject-Specific Analyses in Neuroimaging Data Using ‘Z-Score’ Methods.” J Neurotrauma. 10.1089/neu.2024.0546

Gimbel, S.I., Hungerford, L.D., Twamley, E.W., Ettenhofer, M.L., 2024b. White Matter Organization and Cortical Thickness Differ Among Active Duty Service Members With Chronic Mild, Moderate, and Severe Traumatic Brain Injury. J Neurotrauma 41, 818–835. 10.1089/neu.2023.0336

Harris, N.G., Verley, D.R., Gutman, B.A., Sutton, R.L., 2016. Bi-directional changes in fractional anisotropy after experiment TBI: Disorganization and reorganization? NeuroImage 133, 129–143. 10.1016/j.neuroimage.2016.03.012

Harris, N. G., Verley, D.R., Gutman, B.A., Thompson, P.M., Yeh, H.J., Brown, J.A., 2016. Disconnection and hyper-connectivity underlie reorganization after TBI: A rodent functional connectomic analysis. Exp Neurol 277, 124–138. 10.1016/j.expneurol.2015.12.020

Immonen, R., Harris, N.G., Wright, D., Johnston, L., Manninen, E., Smith, G., Paydar, A., Branch, C., Grohn, O., 2019a. Imaging biomarkers of epileptogenecity after traumatic brain injury – Preclinical frontiers. Neurobiology of Disease 123, 75–85. 10.1016/j.nbd.2018.10.008

Immonen, R., Smith, G., Brady, R.D., Wright, D., Johnston, L., Harris, N.G., Manninen, E., Salo, R., Branch, C., Duncan, D., Cabeen, R., Ndode-Ekane, X.E., Gomez, C.S., Casillas-Espinosa, P.M., Ali, I., Shultz, S.R., Andrade, P., Puhakka, N., Staba, R.J., O’Brien, T.J., Toga, A.W., Pitkänen, A., Gröhn, O., 2019b. Harmonization of pipeline for preclinical multicenter MRI biomarker discovery in a rat model of post-traumatic epileptogenesis. Epilepsy Research 150, 46–57. 10.1016/j.eplepsyres.2019.01.001

Jenkinson, M., Beckmann, C.F., Behrens, T.E.J., Woolrich, M.W., Smith, S.M., 2012. FSL. Neuroimage 62, 782–790. 10.1016/j.neuroimage.2011.09.015

Jeurissen, B., Leemans, A., Tournier, J.-D., Jones, D.K., Sijbers, J., 2013. Investigating the prevalence of complex fiber configurations in white matter tissue with diffusion magnetic resonance imaging. Hum Brain Mapp 34, 2747– 2766. 10.1002/hbm.22099

Jorge, R.E., Acion, L., White, T., Tordesillas-Gutierrez, D., Pierson, R., Crespo-Facorro, B., Magnotta, V.A., 2012. White matter abnormalities in veterans with mild traumatic brain injury. Am J Psychiatry 169, 1284–1291. 10.1176/appi.ajp.2012.12050600

Kellner, E., Dhital, B., Kiselev, V.G., Reisert, M., 2016. Gibbs-ringing artifact removal based on local subvoxel-shifts: Gibbs-Ringing Artifact Removal. Magn. Reson. Med. 76, 1574–1581. 10.1002/mrm.26054

Kim, N., Branch, C.A., Kim, M., Lipton, M.L., 2013. Whole brain approaches for identification of microstructural abnormalities in individual patients: comparison of techniques applied to mild traumatic brain injury. PLoS One 8, e59382. 10.1371/journal.pone.0059382

Kislik, G., Fox, R., Korotcov, A.V., Zhou, J., Febo, M., Moghadas, B., Bibic, A., Zou, Y., Wan, J., Koehler, R.C., Adebayo, T., Burns, M.P., McCabe, J.T., Wang, K.K., Huie, J.R., Ferguson, A.R., Paydar, A., Wanner, I.B., Harris, N.G., Investigators, the T.-N., 2025. Improved Injury Detection Through Harmonizing Multi-Site Neuroimaging Data after Experimental TBI: A Translational Outcomes Project in NeuroTrauma (TOP-NT) Consortium Study. 10.1101/2025.04.15.649026

Lansberg, M.G., Thijs, V.N., O’Brien, M.W., Ali, J.O., de Crespigny, A.J., Tong, D.C., Moseley, M.E., Albers, G.W., 2001. Evolution of apparent diffusion coefficient, diffusion-weighted, and T2-weighted signal intensity of acute stroke. AJNR Am J Neuroradiol 22, 637–644.

LaPlaca, M.C., Huie, J.R., Alam, H.B., Bachstetter, A.D., Bayir, H., Bellgowan, P.F., Cummings, D., Dixon, C.E., Ferguson, A.R., Ferland-Beckham, C., Floyd, C.L., Friess, S.H., Galanopoulou, A.S., Hall, E.D., Harris, N.G., Hawkins, B.E., Hicks, R.R., Hulbert, L.E., Johnson, V.E., Kabitzke, P.A., Lafrenaye, A.D., Lemmon, V.P., Lifshitz, C.W., Lifshitz, J., Loane, D.J., Misquitta, L., Nikolian, V.C., Noble-Haeusslein, L.J., Smith, D.H., Taylor-Burds, C., Umoh, N., Vovk, O., Williams, A.M., Young, M., Zai, L.J., 2021. Pre-Clinical Common Data Elements for Traumatic Brain Injury Research: Progress and Use Cases. Journal of Neurotrauma 38, 1399–1410. 10.1089/neu.2020.7328

Lee, S.-H., Ban, W., Shih, Y.-Y.I., 2020. BrkRaw/bruker: BrkRaw v0.3.3. 10.5281/ZENODO.3877179

Li, L., Chopp, M., Ding, G., Li, Q., Mahmood, A., Jiang, Q., 2017. Chronic global analysis of vascular permeability and cerebral blood flow after bone marrow stromal cell treatment of traumatic brain injury in the rat: A long-term MRI study. Brain Res 1675, 61–70. 10.1016/j.brainres.2017.09.007

Lin, Y., Pan, Y., Wang, M., Huang, X., Yin, Y., Wang, Y., Jia, F., Xiong, W., Zhang, N., Jiang, J., 2012. Blood-brain barrier permeability is positively correlated with cerebral microvascular perfusion in the early fluid percussion-injured brain of the rat. Lab Invest 92, 1623–1634. 10.1038/labinvest.2012.118

Ling, J.M., Peña, A., Yeo, R.A., Merideth, F.L., Klimaj, S., Gasparovic, C., Mayer, A.R., 2012. Biomarkers of increased diffusion anisotropy in semi-acute mild traumatic brain injury: a longitudinal perspective. Brain 135, 1281–1292. 10.1093/brain/aws073

Lipton, M.L., Kim, N., Park, Y.K., Hulkower, M.B., Gardin, T.M., Shifteh, K., Kim, M., Zimmerman, M.E., Lipton, R.B., Branch, C.A., 2012. Robust detection of traumatic axonal injury in individual mild traumatic brain injury patients: intersubject variation, change over time and bidirectional changes in anisotropy. Brain Imaging Behav 6, 329–342. 10.1007/s11682-012-9175-2

Maas, A.I.R., Menon, D.K., David Adelson, P.D., Andelic, N., Bell, M.J., Belli, A., Bragge, P., Brazinova, A., Büki, A., Chesnut, R.M., Citerio, G., Coburn, M., Jamie Cooper, D., Tamara Crowder, A., Czeiter, E., Czosnyka, M., Diaz-Arrastia, R., Dreier, J.P., Duhaime, A.C., Ercole, A., van Essen, T.A., Feigin, V.L., Gao, G., Giacino, J., Gonzalez-Lara, L.E., Gruen, R.L., Gupta, D., Hartings, J.A., Hill, S., Jiang, J.Y., Ketharanathan, N., Kompanje, E.J.O., Lanyon, L., Laureys, S., Lecky, F., Levin, H., Lingsma, H.F., Maegele, M., Majdan, M., Manley, G., Marsteller, J., Mascia, L., McFadyen, C., Mondello, S., Newcombe, V., Palotie, A., Parizel, P.M., Peul, W., Piercy, J., Polinder, S., Puybasset, L., Rasmussen, T.E., Rossaint, R., Smielewski, P., Söderberg, J., Stanworth, S.J., Stein, M.B., von Steinbüchel, N., Stewart, W., Steyerberg, E.W., Stocchetti, N., Synnot, A., Te Ao, B., Tenovuo, O., Theadom, A., Tibboel, D., Videtta, W., Wang, K.K.W., Huw Williams, W., Wilson, L., Yaffe, K., Adams, H., Allanson, J., Coles, J., Hutchinson, P.J., Kolias, A.G., Sahakian, B.J., Stamatakis, E., Williams, G., Agnoletti, V., Martino, C., Masala, A., Teodorani, G., Zumbo, F., Amrein, K., Ezer, E., Kolumbán, B., Kovács, N., Melegh, B., Nyirádi, J., Sorinola, A., Vámos, Z., Andaluz, N., Anke, A., Frisvold, S.K., Antoni, A., van As, A.B., Figaji, A., Audibert, G., Azaševac, A., Dilvesi, D., Golubović, J., Jelača, B., Karan, M., Kolundžija, K., Negru, A., Vulekovic, P., Azouvi, P., Azzolini, M.L., Beretta, L., Baciu, C., Beqiri, V., Chevallard, G., Chieregato, A., Sacchi, M., Badenes, R., Belda, F.J., Bilotta, F., Lozano, A., Barlow, K.M., Schneider, K.J., Bartels, R., den Boogert, H., Hoedemaekers, C., Sir, Ö., Bauerfeind, U., Lefering, R., Schäfer, N., Beauchamp, M., Gravel, J., Beer, D., Beer, R., Helbok, R., Höfer, S., Bellander, B.M., Nelson, D., Bellier, R., Benard, T., Carise, E., Dahyot-Fizelier, C., Giraud, B., Benali, H., Bernard, F., Bertolini, G., Masson, S., Blaabjerg, M., Rosenlund, C., Schou, R.F., Boutis, K., Bouzat, P., Francony, G., Manhes, P., Payen, J.F., Brooks, B., Dewey, D., Emery, C.A., Freedman, S., Kramer, A., Brorsson, C., Koskinen, L.O., Sundström, N., Bullinger, M., Burns, E., Calappi, E., Ortolano, F., Cameron, P., Castaño-León, A.M., Gómez López, P.A., Lagares, A., Causin, F., Freo, U., Persona, P., Rossi, S., Christie, B., Cnossen, M., Dippel, D., Foks, K., Haagsma, J.A., Haitsma, I., Huijben, J.A., van der Jagt, M., Nieboer, D., Volovici, V., Voormolen, D.C., Collett, J., Dawes, H., Esser, P., van Heugten, C., Della Corte, F., Grossi, F., Craig, W., Csato, G., Csomos, A., Curry, N., Dematteo, C., Meade, M., Depreitere, B., van Dijck, J., de Ruiter, G.C.W., Vleggeert-Lankamp, C., Dizdarevic, K., Donoghue, E., Gantner, D., Murray, L., Trapani, T., Vallance, S., Duek, O., Lazar, I., Dulière, G.L., Maréchal, H., Dzeko, A., Eapen, G., Jankowski, S., English, S., Fergusson, D., Osmond, M., Fabricius, M., Kondziella, D., Feng, J., Hui, J., Fleming, J., Latini, R., Gagnon, I., Ptito, A., Galanaud, D., Glocker, B., Kamnitsas, K., Ledig, C., Rueckert, D., Gordon, W.A., Gradisek, P., Griesdale, D., Håberg, A.K., van Hecke, W., Smeets, D., Verheyden, J., Vyvere, T.V., Helseth, E., Røe, C., Røise, O., Horton, L., Jacobs, B., van der Naalt, J., Janssens, K., De Keyser, V., Menovsky, T., Van Praag, D., Jones, K.M., Kapš, R., Katila, A., Posti, J., Takala, R., Kaukonen, K.M., Kivisaari, R., Piippo-Karjalainen, A., Raj, R., Tanskanen, P., Kutsogiannis, D., Kyprianou, T., Lamontagne, F., Lauzier, F., Moore, L., Turgeon, A., Legrand, V., Levi, L., Zaaroor, M., Lightfoot, R., Macdonald, S., Major, S., Vajkoczy, P., Wessels, L., Winkler, M.K.L., Wolf, S., Manara, A., Thomas, M., Mattern, J., Sakowitz, O., Vogt, L., Younsi, A., McFadyen, B., McMahon, C., Correia, M.M., Morganti-Kossmann, M.C., Rosenfeld, J.V., Muehlan, H., Schmidt, S., Mukherjee, P., Noirhomme, Q., Oddo, M., Okonkwo, D.O., Oldenbeuving, A.W., Roks, G., Schoonman, G.G., Perlbarg, V., Pichon, N., Pili-Floury, S., Pirinen, M., Pleş, H., Poca, M.A., Radoi, A., Sahuquillo, J., Ragauskas, A., Rocka, S., Real, R.G.L., Telgmann, R., Reed, N., Rhodes, J., Robertson, C., Rosand, J., Rosenthal, G., Salvato, G., Sánchez-Porras, R., Sándor, J., Sangha, G., Schnyer, D., Schöhl, H., Skandsen, T., Stevanovic, A., van Waesberghe, J.V., Stevens, R.D., Taccone, F.S., Taylor, M.S., Zelinkova, V., Temkin, N., Tolias, C.M., Valadka, A.B., Valeinis, E., Vargiolu, A., Vega, E., Vik, A., Vilcinis, R., Wildschut, E., Wood, G., Xirouchaki, N., Zemek, R., 2017. Traumatic brain injury: Integrated approaches to improve prevention, clinical care, and research. Lancet Neurol 16, 987–1048. 10.1016/S1474-4422(17)30371-X

Mayer, A.R., Bedrick, E.J., Ling, J.M., Toulouse, T., Dodd, A., 2014. Methods for identifying subject-specific abnormalities in neuroimaging data. Hum Brain Mapp 35, 5457–5470. 10.1002/hbm.22563

Mayer, A.R., Dodd, A.B., Ling, J.M., Bedrick, E.J., 2024. Revisiting Subject-Specific Analyses in Neuroimaging Data Using “Z-Score” Methods. Journal of Neurotrauma. 10.1089/neu.2024.0434

Mayer, A.R., Ling, J.M., Yang, Z., Pena, A., Yeo, R.A., Klimaj, S., 2012. Diffusion abnormalities in pediatric mild traumatic brain injury. J Neurosci 32, 17961–17969. 10.1523/JNEUROSCI.3379-12.2012

Obenaus, A., Robbins, M., Blanco, G., Galloway, N.R., Snissarenko, E., Gillard, E., Lee, S., Currás-Collazo, M., 2007. Multi-Modal Magnetic Resonance Imaging Alterations in Two Rat Models of Mild Neurotrauma. Journal of Neurotrauma 24, 1147–1160. 10.1089/neu.2006.0211

Prinz, F., Schlange, T., Asadullah, K., 2011. Believe it or not: how much can we rely on published data on potential drug targets? Nat Rev Drug Discov 10, 712–712. 10.1038/nrd3439-c1

Radabaugh, H.L., Harris, N.G., Wanner, I.B., Burns, M.P., McCabe, J.T., Korotcov, A.V., Dardzinski, B.J., Zhou, J., Koehler, R.C., Wan, J., Allende Labastida, J., Moghadas, B., Bibic, A., Febo, M., Kobeissy, F.H., Zhu, J., Rubenstein, R., Hou, J., Bose, P.K., Apiliogullari, S., Beattie, M.S., Bresnahan, J.C., Rosi, S., Huie, J.R., Ferguson, A.R., Wang, K.K.W., the TOP-NT Investigators, 2025. Translational Outcomes Project in Neurotrauma (TOP-NT) Pre-Clinical Consortium Study: A Synopsis. Journal of Neurotrauma neu.2023.0654. 10.1089/neu.2023.0654

Rubovitch, V., Ten-Bosch, M., Zohar, O., Harrison, C.R., Tempel-Brami, C., Stein, E., Hoffer, B.J., Balaban, C.D., Schreiber, S., Chiu, W.-T., Pick, C.G., 2011. A mouse model of blast-induced mild traumatic brain injury. Experimental Neurology 232, 280–289. 10.1016/j.expneurol.2011.09.018

Santana-Gomez, C., Smith, G., Mousavi, A., Shamas, M., Harris, N.G., Staba, R., 2024. The Surgical Method of Craniectomy Differentially Affects Acute Seizures, Brain Deformation, and Behavior in a Traumatic Brain Injury Animal Model. Neurotrauma Rep 5, 969–981. 10.1089/neur.2024.0064

Schneider, G., Fries, P., Wagner-Jochem, D., Thome, D., Laurer, H., Kramann, B., Mautes, A., Hagen, T., 2002. Pathophysiological changes after traumatic brain injury: comparison of two experimental animal models by means of MRI. MAGMA 14, 233–241. 10.1007/BF02668217

Sert, N.P. du, Hurst, V., Ahluwalia, A., Alam, S., Avey, M.T., Baker, M., Browne, W.J., Clark, A., Cuthill, I.C., Dirnagl, U., Emerson, M., Garner, P., Holgate, S.T., Howells, D.W., Karp, N.A., Lazic, S.E., Lidster, K., MacCallum, C.J., Macleod, M., Pearl, E.J., Petersen, O.H., Rawle, F., Reynolds, P., Rooney, K., Sena, E.S., Silberberg, S.D., Steckler, T., Würbel, H., 2020. The ARRIVE guidelines 2.0: Updated guidelines for reporting animal research. PLOS Biology 18, e3000410. 10.1371/journal.pbio.3000410

Smith, D.H., Hicks, R.R., Johnson, V.E., Bergstrom, D.A., Cummings, D.M., Noble, L.J., Hovda, D., Whalen, M., Ahlers, S.T., LaPlaca, M., Tortella, F.C., Duhaime, A.-C., Dixon, C.E., 2015. Pre-Clinical Traumatic Brain Injury Common Data Elements: Toward a Common Language Across Laboratories. J Neurotrauma 32, 1725–1735. 10.1089/neu.2014.3861

Smith, G., Thapak, P., Paydar, A., Ying, Z., Gomez-Pinilla, F., Harris, N., G., 2023. Altering the Trajectory of Perfusion-Diffusion Deficits Using A BDNF Mimetic Acutely After TBI is Associated with Improved Functional Connectivity. PNEURO 1–12. 10.60124/j.PNEURO.2023.10.07

Smith, S.M., 2002. Fast robust automated brain extraction. Hum Brain Mapp 17, 143–155. 10.1002/hbm.10062

Smith, S.M., Jenkinson, M., Woolrich, M.W., Beckmann, C.F., Behrens, T.E.J., Johansen-Berg, H., Bannister, P.R., De Luca, M., Drobnjak, I., Flitney, D.E., Niazy, R.K., Saunders, J., Vickers, J., Zhang, Y., De Stefano, N., Brady, J.M., Matthews, P.M., 2004. Advances in functional and structural MR image analysis and implementation as FSL. NeuroImage, Mathematics in Brain Imaging 23, S208–S219. 10.1016/j.neuroimage.2004.07.051

Stein, D.G., 2015. Embracing failure: What the Phase III progesterone studies can teach about TBI clinical trials. Brain Inj 29, 1259–1272. 10.3109/02699052.2015.1065344

Tournier, J.-D., Smith, R., Raffelt, D., Tabbara, R., Dhollander, T., Pietsch, M., Christiaens, D., Jeurissen, B., Yeh, C.-H., Connelly, A., 2019. MRtrix3: A fast, flexible and open software framework for medical image processing and visualisation. NeuroImage 202, 116137. 10.1016/j.neuroimage.2019.116137

Trouble at the lab, Oct 18^th^ 2013. The Economist.

Valdés-hernández, P.A., Sumiyoshi, A., Nonaka, H., Haga, R., Ogawa, T., Iturria-medina, Y., Riera, J.J., Kawashima, R., Roche, A., Medical, S., 2011. An in vivo MRI template set for morphometry, tissue segmentation, and fMRI localization in rats 5, 1–19. 10.3389/fninf.2011.00026

Veraart, J., Fieremans, E., Novikov, D.S., 2016. Diffusion MRI noise mapping using random matrix theory. Magnetic Resonance in Medicine 76, 1582–1593. 10.1002/mrm.26059

Wanner, I.-B., McCabe, J.T., Huie, J.R., Harris, N.G., Paydar, A., McMann-Chapman, C., Tobar, A., Korotcov, A., Burns, M.P., Koehler, R.C., Wan, J., Allende Labastida, J., Tong, J., Zhou, J., Davis, L.M., Radabaugh, H.L., Ferguson, A.R., Van Meter, T.E., Febo, M., Bose, P., Wang, K.K., Kobeissy, F., Apiliogullari, S., Zhu, J., Rubenstein, R., Awwad, H.O., and the TOP-NT Consortium Investigators, 2025. Prospective Harmonization, Common Data Elements, and Sharing Strategies for Multicenter Pre-Clinical Traumatic Brain Injury Research in the Translational Outcomes Project in Neurotrauma Consortium. Journal of Neurotrauma. 10.1089/neu.2023.0653

Watts, R., Thomas, A., Filippi, C.G., Nickerson, J.P., Freeman, K., 2014. Potholes and molehills: bias in the diagnostic performance of diffusion-tensor imaging in concussion. Radiology 272, 217–223. 10.1148/radiol.14131856

Woolrich, M.W., Jbabdi, S., Patenaude, B., Chappell, M., Makni, S., Behrens, T., Beckmann, C., Jenkinson, M., Smith, S.M., 2009. Bayesian analysis of neuroimaging data in FSL. Neuroimage 45, S173–186. 10.1016/j.neuroimage.2008.10.055

Wright, D.K., Johnston, L.A., Kershaw, J., Ordidge, R., O’Brien, T.J., Shultz, S.R., 2017. Changes in Apparent Fiber Density and Track-Weighted Imaging Metrics in White Matter following Experimental Traumatic Brain Injury. J Neurotrauma 34, 2109–2118. 10.1089/neu.2016.4730

Zhang, H., Yushkevich, P.A., Alexander, D.C., Gee, J.C., 2006. Deformable registration of diffusion tensor MR images with explicit orientation optimization. Medical Image Analysis, The Eighth International Conference on Medical Imaging and Computer Assisted Intervention – MICCAI 2005 10, 764–785. 10.1016/j.media.2006.06.004

Zvenigorodsky, V., Gruenbaum, B.F., Shelef, I., Horev, A., Azab, A.N., Oleshko, A., Abu-Rabia, M., Negev, S., Zlotnik, A., Melamed, I., Boyko, M., 2024. Evaluation of Blood-Brain Barrier Disruption Using Low- and High-Molecular-Weight Complexes in a Single Brain Sample in a Rat Traumatic Brain Injury Model: Comparison to an Established Magnetic Resonance Imaging Technique. Int J Mol Sci 25, 11241. 10.3390/ijms252011241

