## Supplemental for "Unbiased Population-Based Statistics to Obtain Pathologic Burden of Injury after Experimental TBI"

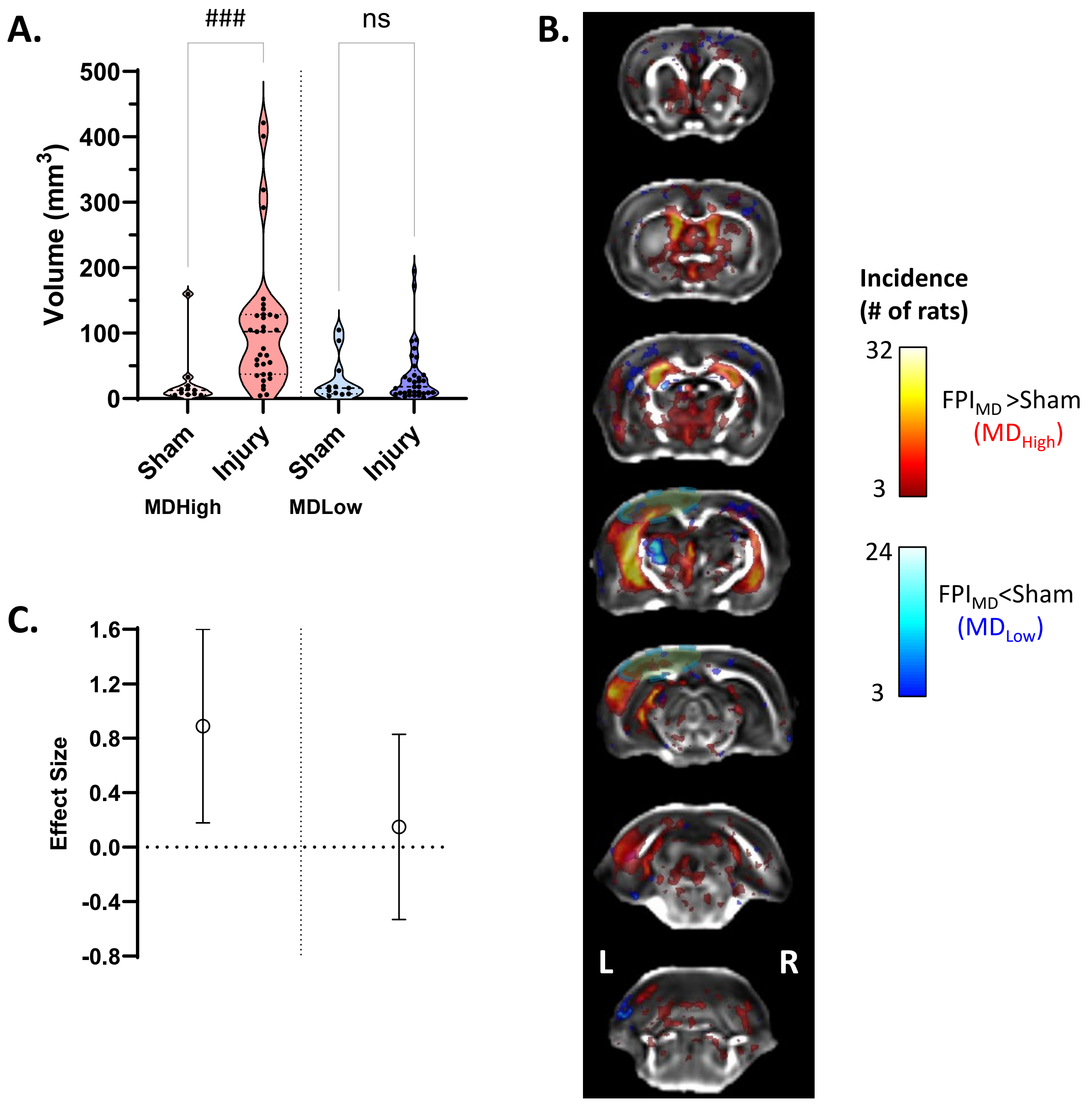
A.
###
ns
Volume (mm3)
500
400
300
200
100
0
Sham
Injury
Sham
Injury
MDHigh
MDLow
B.
L
R
Incidence
(# of rats)
32
3
FPIMD >Sham
(MDHigh)
24
3
FPIMD<Sham
(MDLow)
C.
Effect Size
1.6
1.2
0.8
0.4
0.0
-0.4
-0.8

MD Sup1

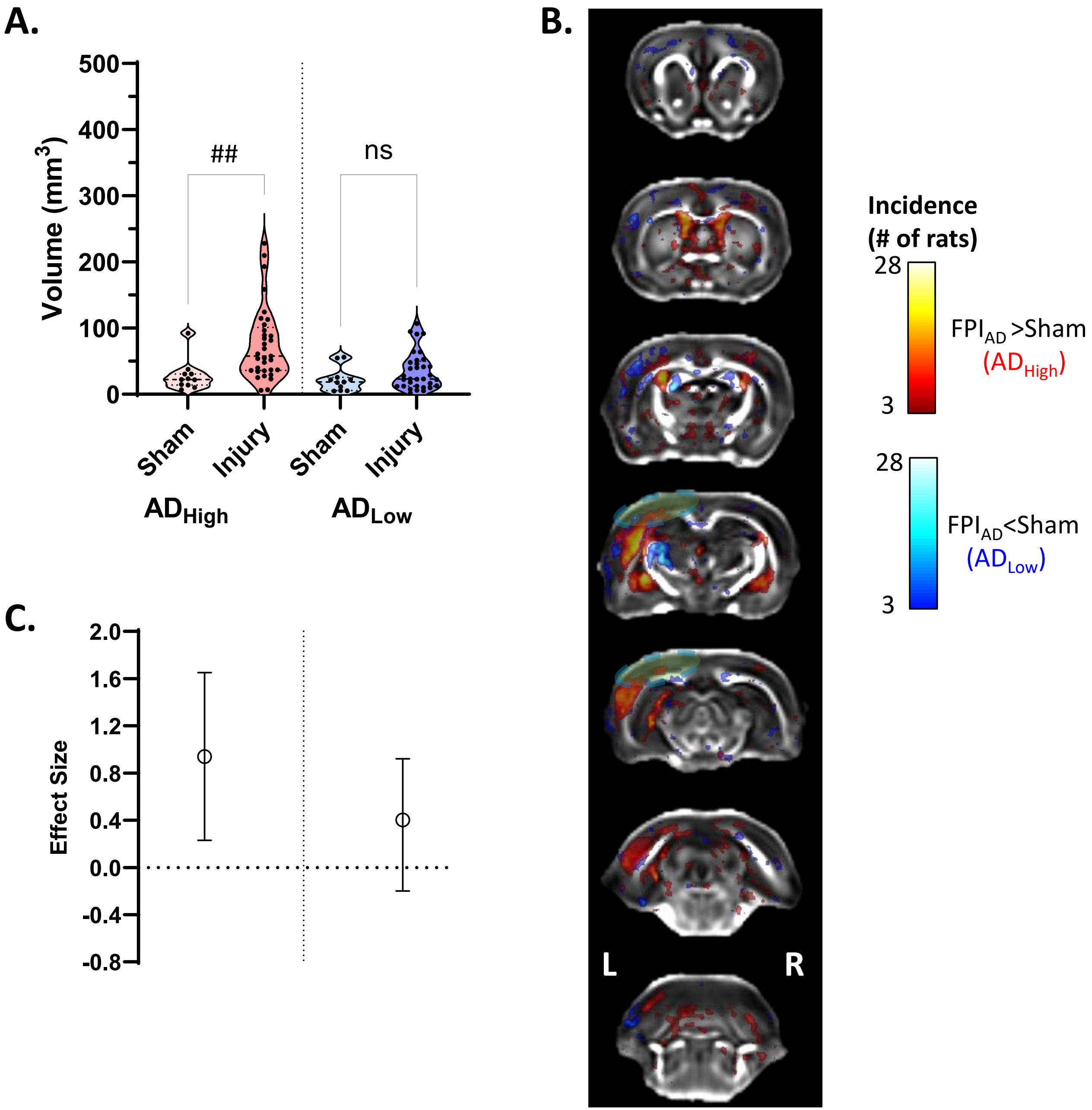
A.
Volume (mm3)
500
400
300
200
100
0
##
ns
Sham
Injury
Sham
Injury
ADHigh
ADLow
B.
L
R
Incidence
(# of rats)
28
3
FPIAD >Sham
(ADHigh)
28
3
FPIAD<Sham
(ADLow)
C.
Effect Size
2.0
1.6
1.2
0.8
0.4
0.0
-0.4
-0.8

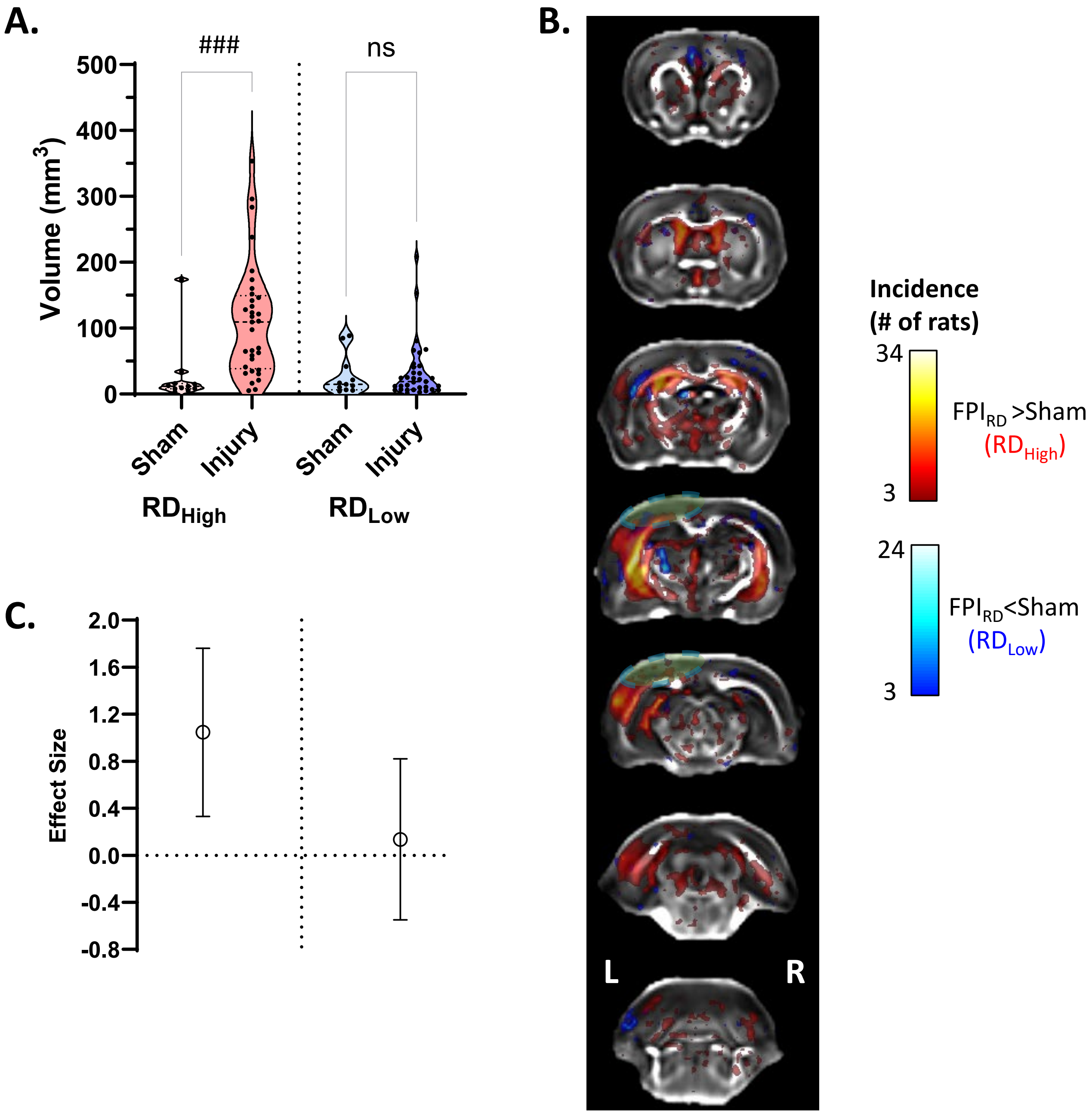
A.
###
ns
Volume (mm3)
500
400
300
200
100
0
Sham
Injury
Sham
Injury
RDHigh
RDLow
B.
Incidence
(# of rats)
34
3
FPIRD >Sham
(RDHigh)
24
3
FPIRD<Sham
(RDLow)
L
R
C.
Effect Size
2.0
1.6
1.2
0.8
0.4
0.0
-0.4
-0.8

RD Sup3

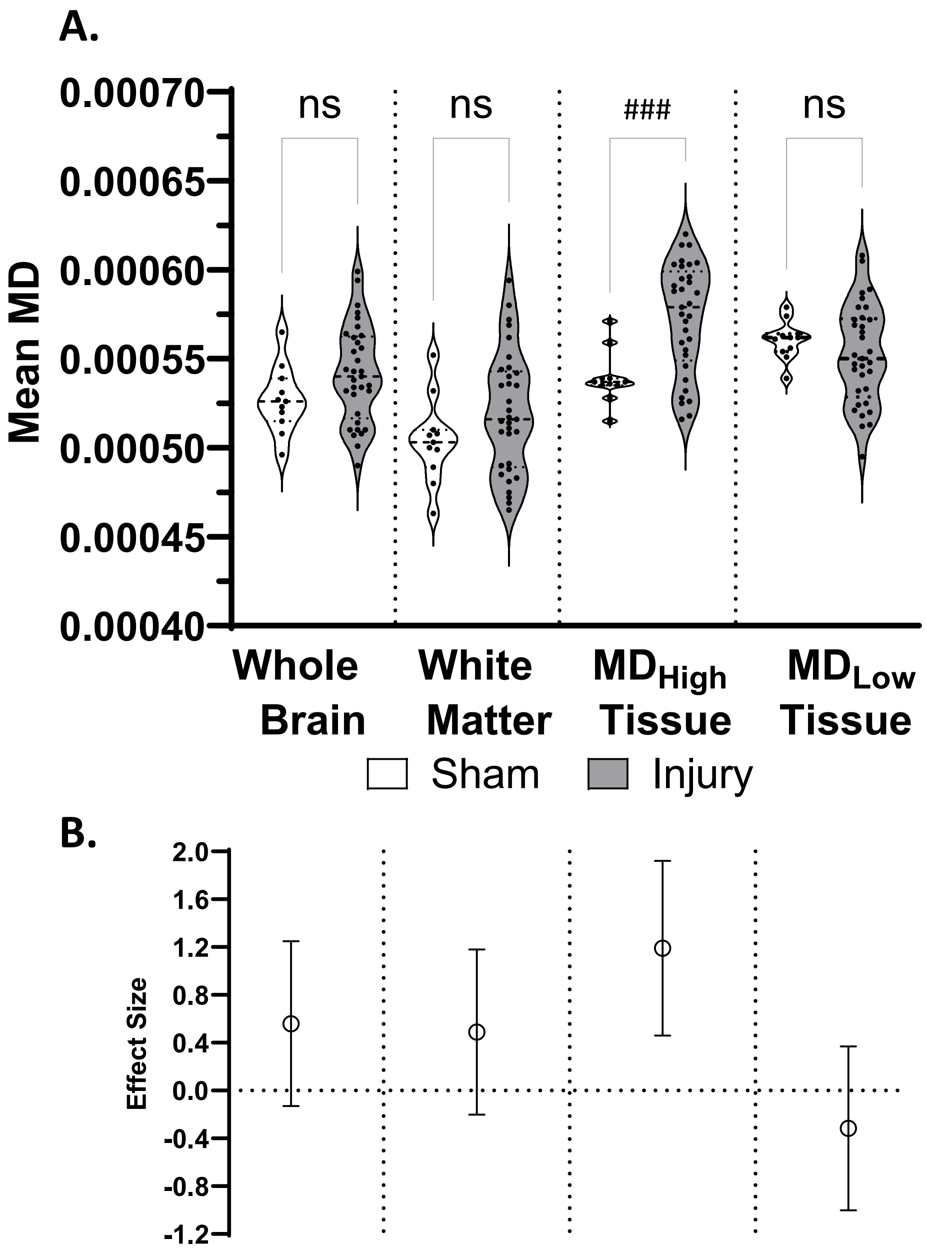
A.
0.00070
0.00065
0.00060
0.00055
0.00050
0.00045
0.00040
Mean MD
ns
ns
###
ns
Whole Brain
White Matter
MD High Tissue
MD Low Tissue
Sham
Injury
B.
2.0
1.6
1.2
0.8
0.4
0.0
-0.4
-0.8
-1.2
Effect Size

MD Sup4

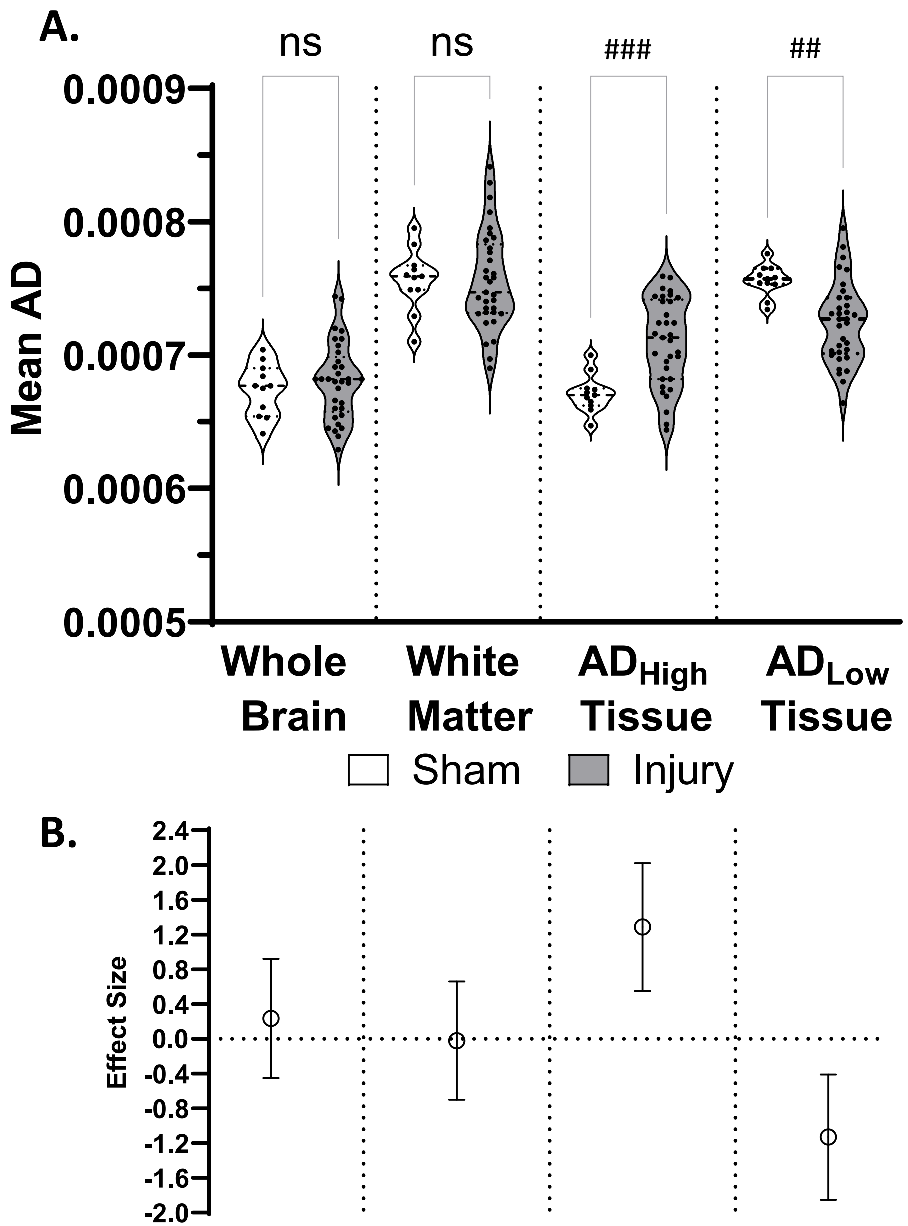
A.
ns
ns
###
##
0.0009
0.0008
0.0007
0.0006
0.0005
Mean AD
Whole Brain
White Matter
AD High Tissue
AD Low Tissue
Sham
Injury
B.
2.4
2.0
1.6
1.2
0.8
0.4
0.0
-0.4
-0.8
-1.2
-1.6
-2.0
Effect Size

AD Sup5

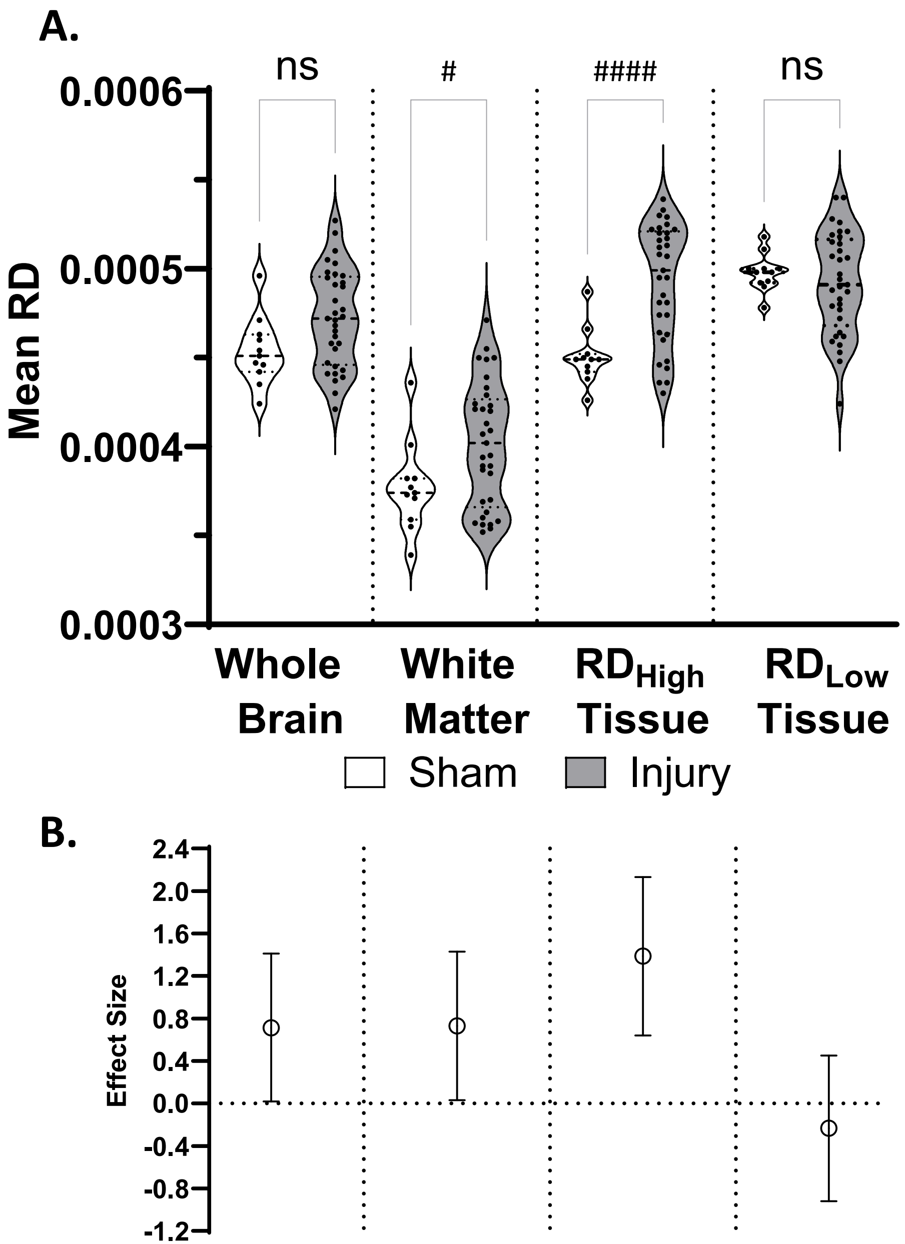
A.
ns
#
####
ns
0.0006
0.0005
0.0004
0.0003
Mean RD
Whole Brain
White Matter
RD High Tissue
RD Low Tissue
Sham
Injury
B.
2.4
2.0
1.6
1.2
0.8
0.4
0.0
-0.4
-0.8
-1.2
Effect Size

RD Sup6

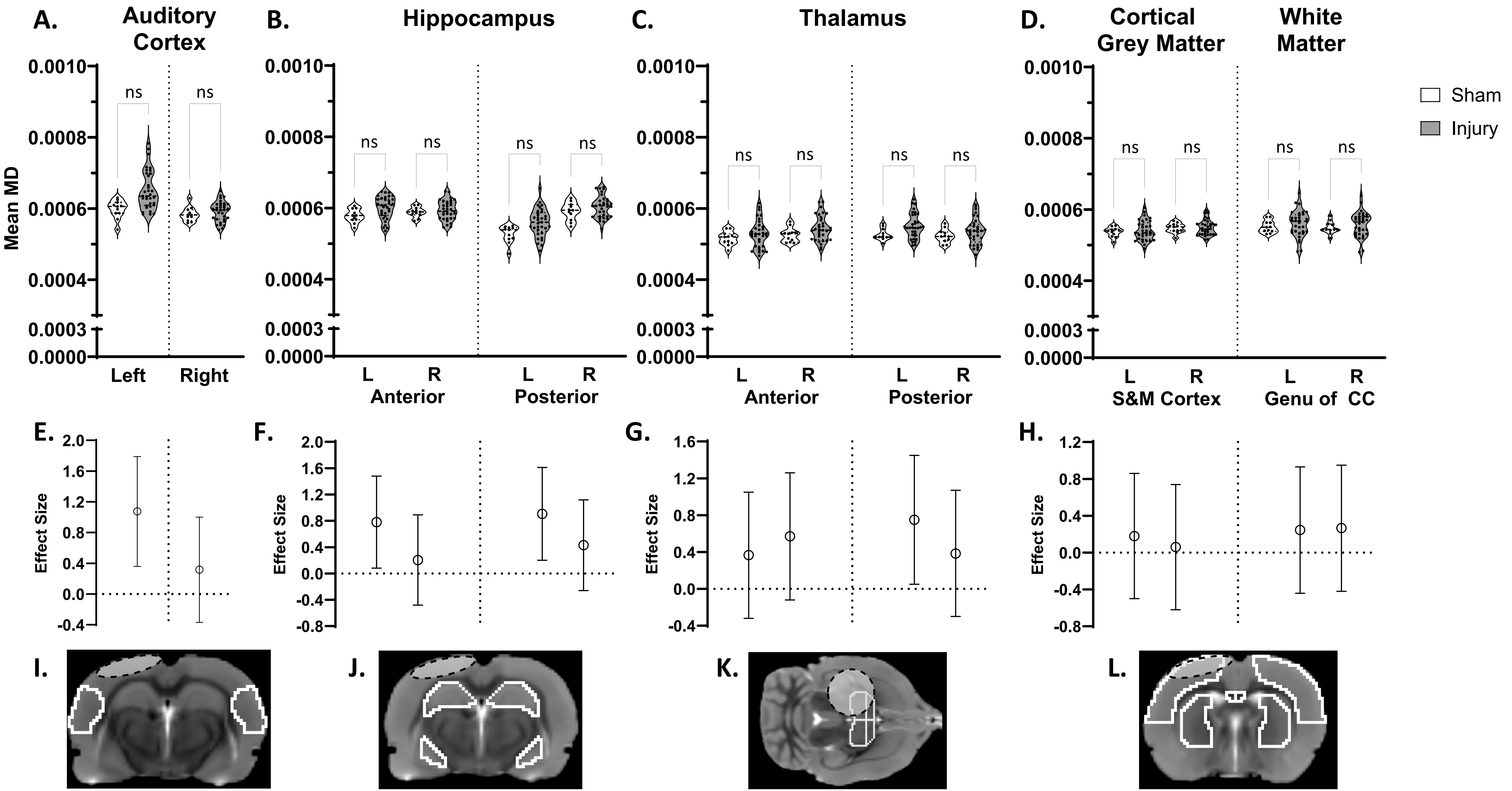
A.
Auditory Cortex
B.
Hippocampus
C.
Thalamus
D.
Cortical Grey Matter
White Matter
Sham
Injury
Mean MD
0.0010
0.0008
0.0006
0.0004
0.0003
0.0000
ns
Left
Right
L
R
Anterior
Posterior
S&M Cortex
Genu of CC
E.
F.
G.
H.
Effect Size
I.
J.
K.
L.

MD Sup7

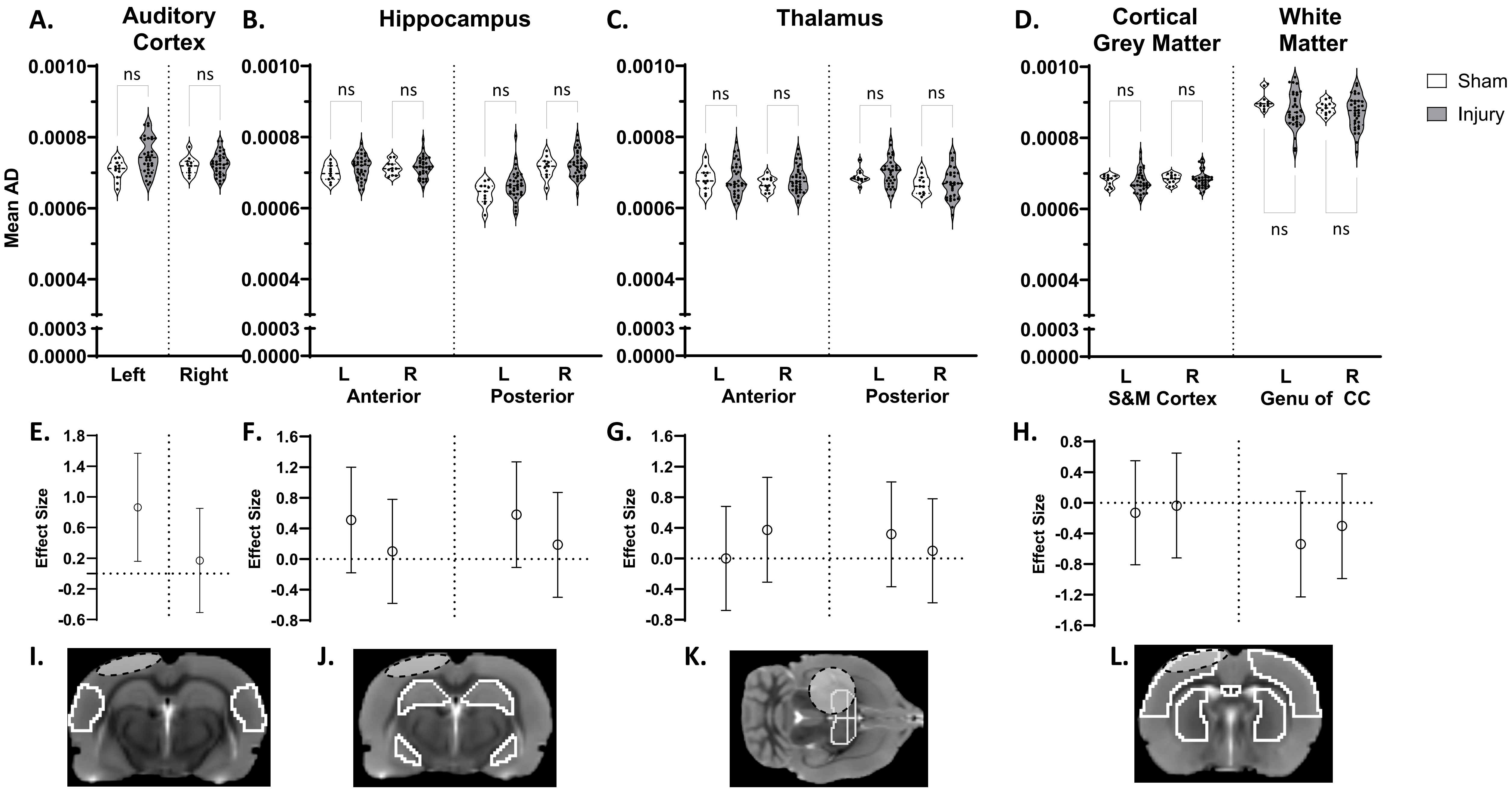
A.
Auditory Cortex
B.
Hippocampus
C.
Thalamus
D.
Cortical Grey Matter
White Matter
Sham
Injury
Mean AD
0.0010
0.0008
0.0006
0.0004
0.0003
0.0000
ns
Left
Right
L
R
Anterior
Posterior
S&M Cortex
Genu of CC
E.
F.
G.
H.
Effect Size
1.8
1.4
1.6
1.2
1.0
0.8
0.6
0.4
0.2
0.0
-0.2
-0.4
-0.6
-0.8
-1.2
-1.6
I.
J.
K.
L.

AD Sup8

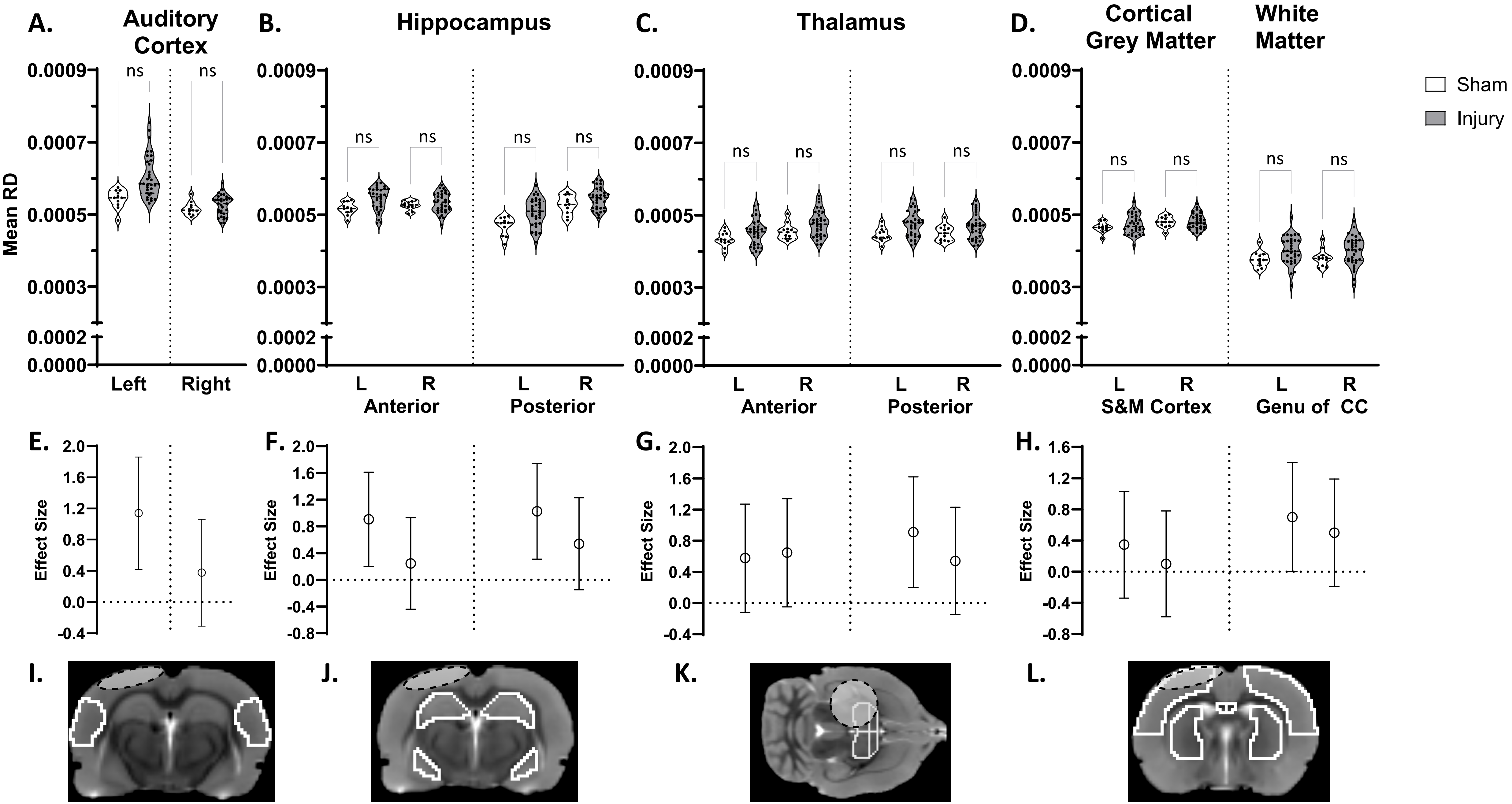
A.
Auditory Cortex
B.
Hippocampus
C.
Thalamus
D.
Cortical Grey Matter
White Matter
Sham
Injury
Mean RD
0.0009
0.0007
0.0005
0.0003
0.0002
0.0000
ns
Left
Right
L
R
Anterior
Posterior
S&M Cortex
Genu of CC
E.
F.
G.
H.
Effect Size
2.0
1.6
1.2
0.8
0.4
0.0
-0.4
-0.8
I.
J.
K.
L.

RD Sup9

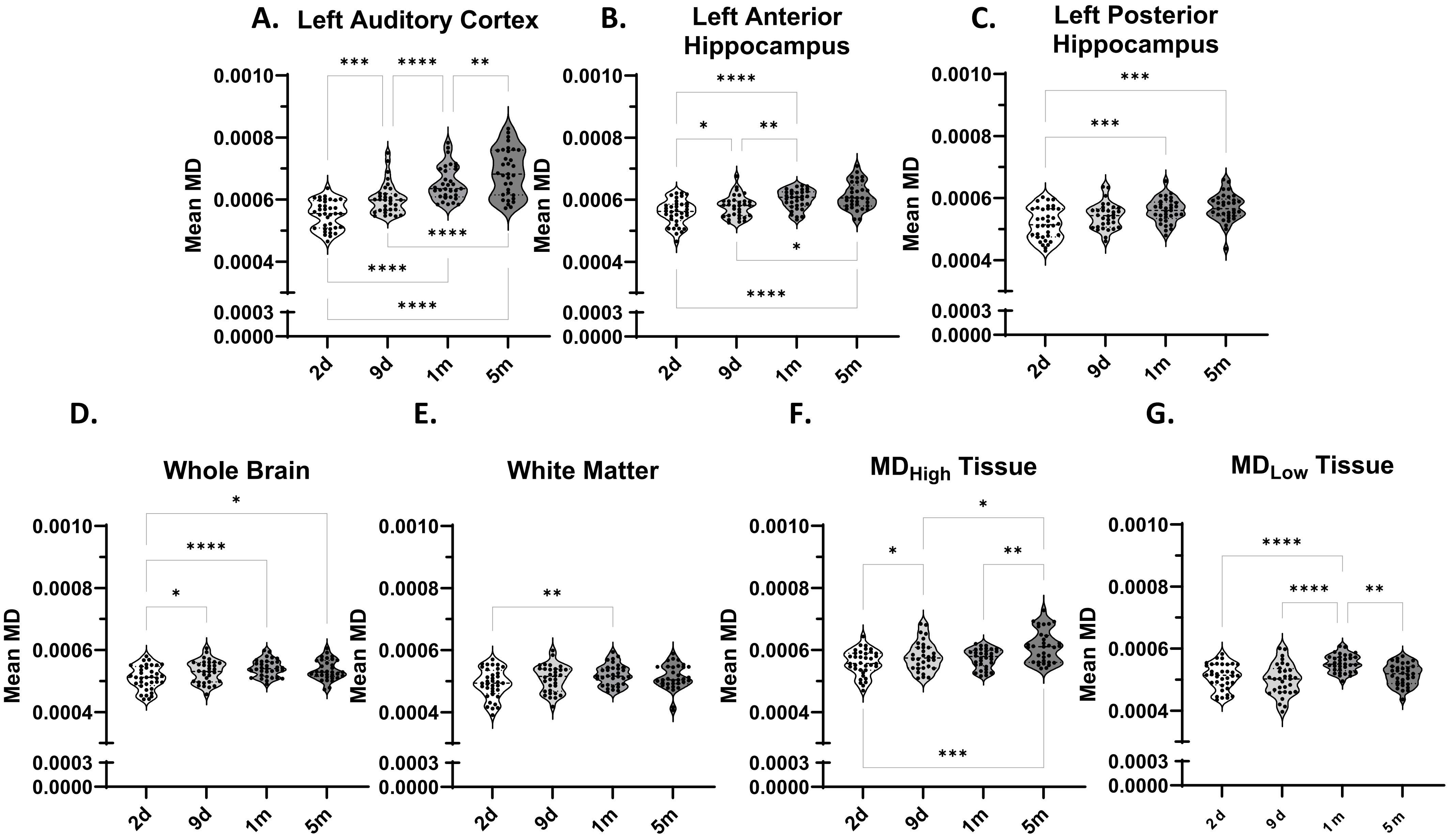
A. Left Auditory Cortex
B. Left Anterior Hippocampus
C. Left Posterior Hippocampus
D.
Whole Brain
E.
White Matter
F.
MD$_{High}$ Tissue
G.
MD$_{Low}$ Tissue
Mean MD
0.0010
0.0008
0.0006
0.0004
0.0003
0.0000
2d
9d
1m
5m
*
**
***
****

MD Sup10

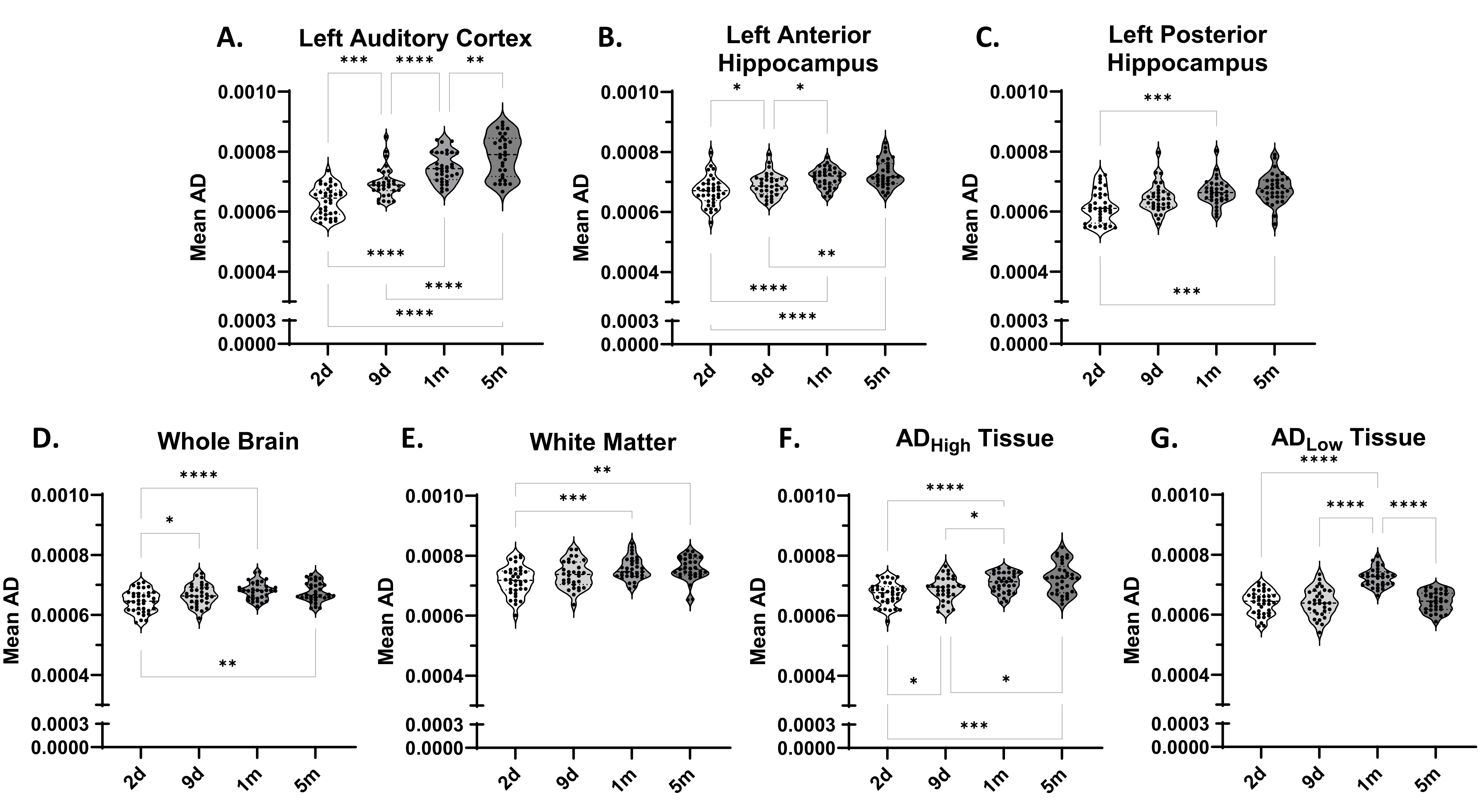
A.
Left Auditory Cortex
B.
Left Anterior Hippocampus
C.
Left Posterior Hippocampus
D.
Whole Brain
E.
White Matter
F.
$AD_{High}$ Tissue
G.
$AD_{Low}$ Tissue
Mean AD
0.0010
0.0008
0.0006
0.0004
0.0003
0.0000
2d
9d
1m
5m
*
**
***
****

AD Sup11

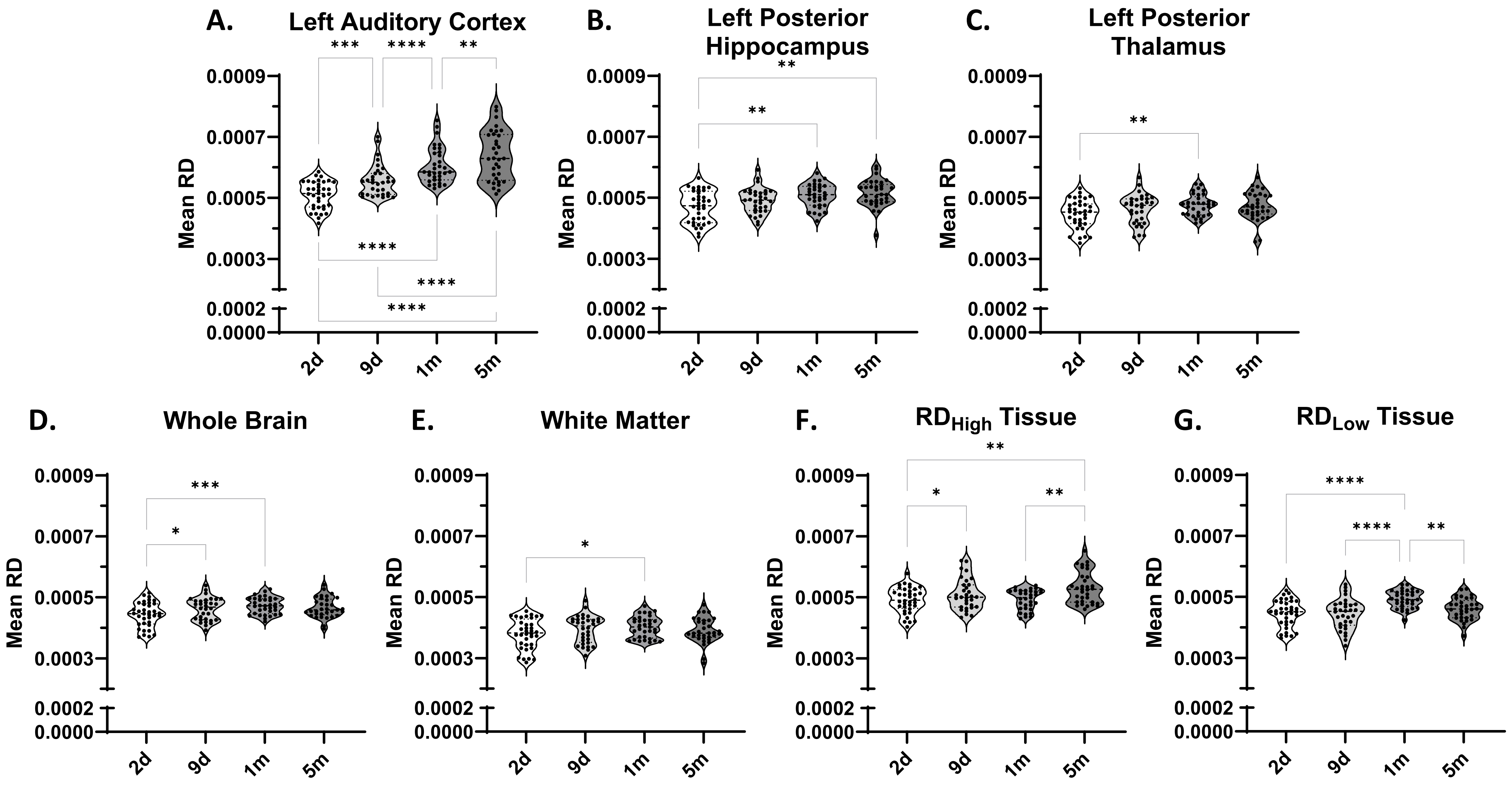
A.
Left Auditory Cortex
B.
Left Posterior Hippocampus
C.
Left Posterior Thalamus
D.
Whole Brain
E.
White Matter
F.
RDHigh Tissue
G.
RDLow Tissue
Mean RD
0.0009
0.0007
0.0005
0.0003
0.0002
0.0000
2d
9d
1m
5m
*
**
***
****

RD Sup12

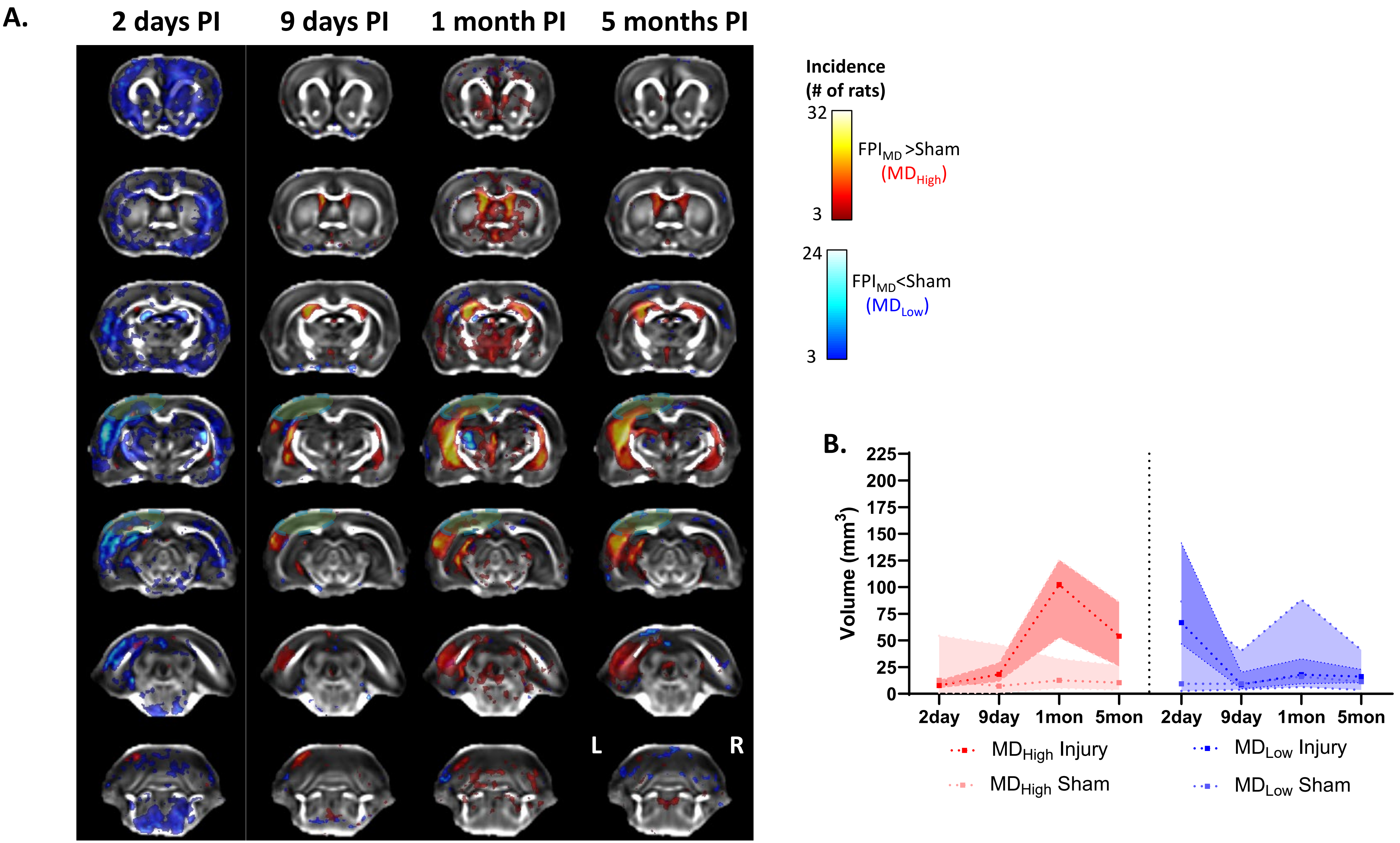

A.
2 days PI
9 days PI
1 month PI
5 months PI
L
R
Incidence
(# of rats)
32
3
FPI$_{MD}$ >Sham
(MD$_{High}$)
24
3
FPI$_{MD}$<Sham
(MD$_{Low}$)
B.
Volume (mm$^3$)
225
200
175
150
125
100
75
50
25
0
2day
9day
1mon
5mon
2day
9day
1mon
5mon
MD$_{High}$ Injury
MD$_{High}$ Sham
MD$_{Low}$ Injury
MD$_{Low}$ Sham


MD Sup13

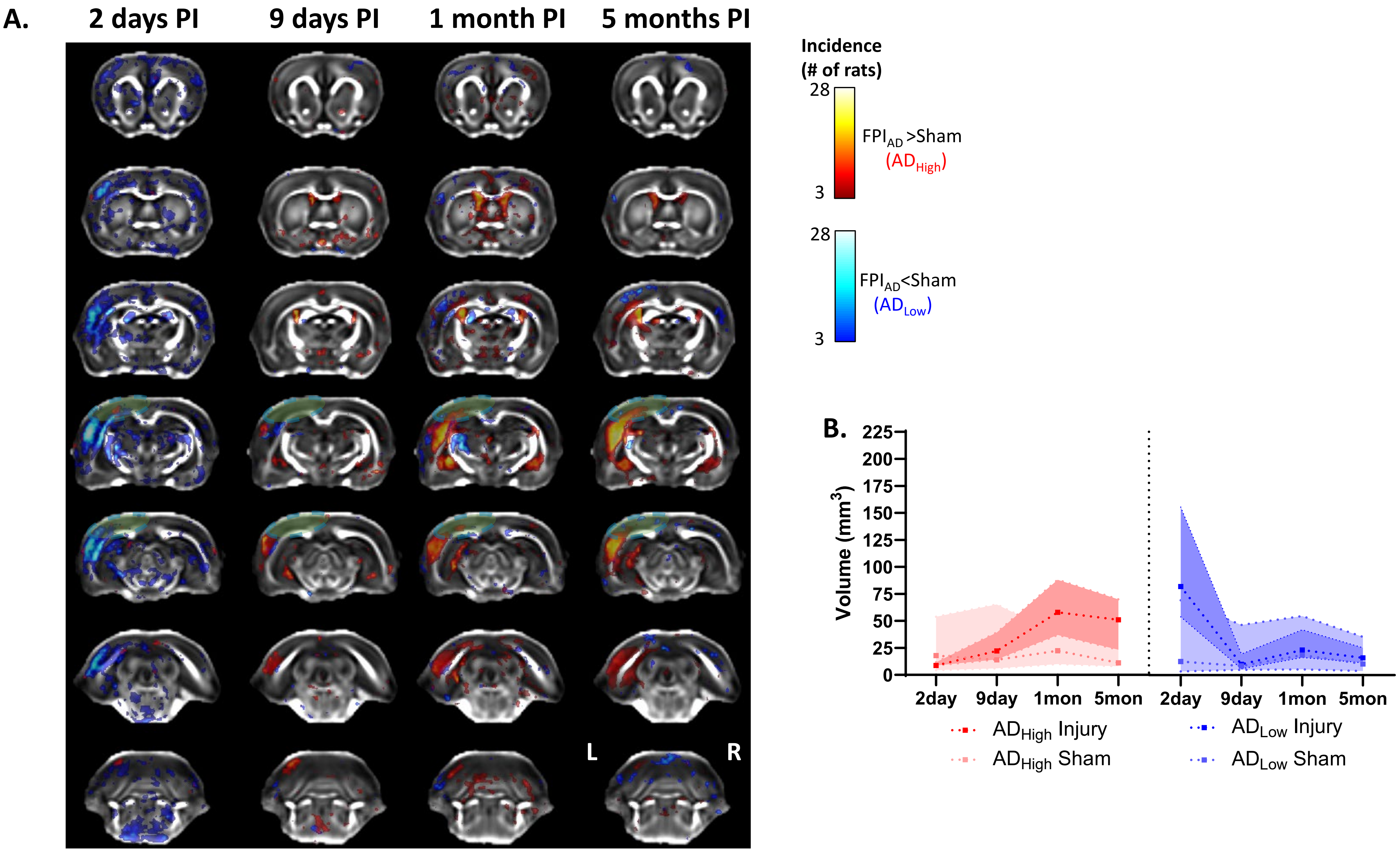

A.
2 days PI
9 days PI
1 month PI
5 months PI
L
R
Incidence
(# of rats)
28
3
FPI$_{AD}$ >Sham
(AD$_{High}$)
28
3
FPI$_{AD}$<Sham
(AD$_{Low}$)
B.
Volume (mm$^3$)
225
200
175
150
125
100
75
50
25
0
2day
9day
1mon
5mon
2day
9day
1mon
5mon
AD$_{High}$ Injury
AD$_{High}$ Sham
AD$_{Low}$ Injury
AD$_{Low}$ Sham


AD Sup14

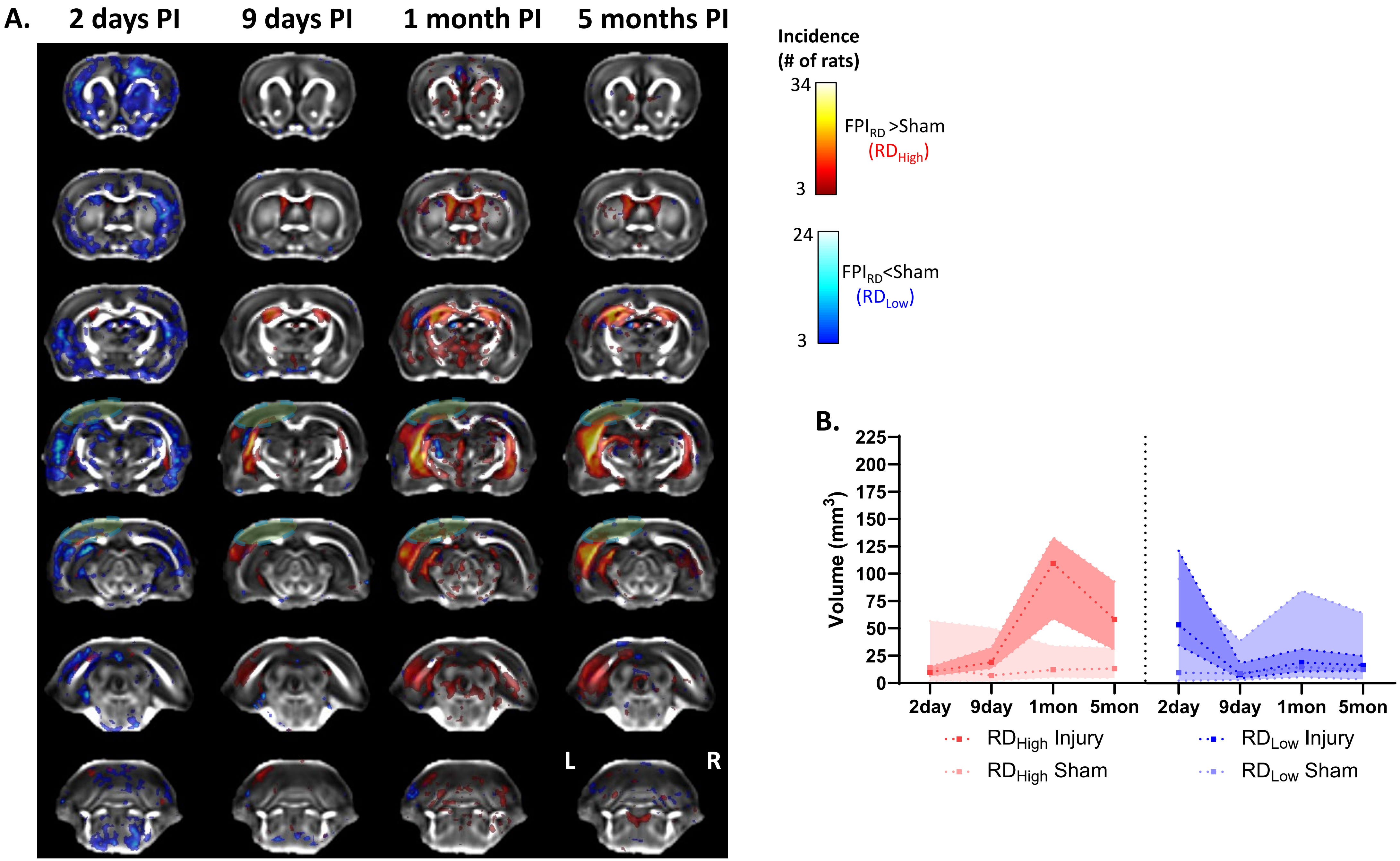
A.
2 days PI
9 days PI
1 month PI
5 months PI
L
R
Incidence
(# of rats)
34
3
FPI$_{RD}$ >Sham
(RD$_{High}$)
24
3
FPI$_{RD}$<Sham
(RD$_{Low}$)
B.
Volume (mm$^3$)
225
200
175
150
125
100
75
50
25
0
2day
9day
1mon
5mon
2day
9day
1mon
5mon
RD$_{High}$ Injury
RD$_{High}$ Sham
RD$_{Low}$ Injury
RD$_{Low}$ Sham

RD Sup15
